# Abstract and Concrete Working Memory Information in Human Visual Cortex

**DOI:** 10.64898/2026.05.23.727368

**Authors:** Or Yizhar, Fabio Bauer, Inés Pont Sanchis, Felix Bröhl, Bernhard J. Spitzer

## Abstract

Neural correlates of visuospatial working memory (WM) information have been found in frontal, parietal, and early visual areas, but the principles by which distributed WM storage is topographically organised remain subject to debate. Here, we used functional magnetic resonance imaging, representational geometry analysis, and vision models to examine the extent to which WM representations across the cortical hierarchy may differ in terms of visuospatial abstraction. During retro-cued WM maintenance of rotated real-world objects, we found robust encoding of the objects’ orientation in both parietal and occipital visual areas. The representational format of this encoding was independent of the objects’ physical appearance and was surprisingly invariant across areas, indicating a high level of visuospatial abstraction even in the early visual cortex. Interestingly, we also found evidence for an orthogonal, object-specific orientation representation within the same areas, likely reflecting the sample stimuli’s concrete visual appearance. The latter—but not the former—type of distributed representation emerged also in a contemporary vision model. Together, the findings indicate the widespread co-existence of abstract (generalised) and concrete-visual representations multiplexed within the same brain areas. In contrast, prototypical orientation biases emerged only in the parietal cortex, suggesting a distinction between generalisation and categorisation in WM abstraction.

## Introduction

The ability to temporarily hold and manipulate information for upcoming tasks, known as working memory (WM), is essential for virtually every form of higher cognition. WM function has traditionally been attributed to the prefrontal cortex^1,2^ but is also assumed to recruit distributed brain areas^3^, including parietal and sensory cortices^4–6^. Multivariate pattern analysis of human neuroimaging data has routinely read out visual WM contents from the early visual cortex^7–10^, as well as from extrastriate^11,12^ and parietal areas^13–15^. The distinct roles of these brain areas in WM storage and the principles that govern them remain a matter of debate.

According to one view, distributed WM storage is topographically organised according to the information’s levels of abstraction, where early sensory areas hold concrete (e.g., visual) details, whereas more anterior areas retain more generalised or categorical representations^16–19^. In line with this view, the EVC was found to encode memorised stimulus features such as grating orientations in activity patterns similar to those observed during perception^7–10,10,20–23^ (but see ^24–28^ for evidence that mnemonic and perceptual neural formats may differ), whilst extrastriate and parietal areas appear to encode visual WM information in a more categorical fashion^17,29,30^. Most anteriorly, the prefrontal cortex maintains task rules^31,32^, abstract magnitude^33–35^ and semantic categories^36^. Goal-directed abstraction in higher-level areas may reduce the amount of information to be stored and render mnemonic representations more resistant to perceptual interference^12,13,37,38^.

Recently, however, evidence has emerged that even early sensory areas might maintain WM information in more abstract formats than previously thought^8,29,39^. For instance, a fMRI study by Kwak and Curtis (2022) showed that WM representations of grating orientation in EVC shared neural formats with those of motion direction in the same area, indicating a degree of generalisation or abstraction from visually dissimilar inputs. Specifically, the results indicated that during WM processing, EVC may represent a surrogate imagery stimulus, a “line” oriented in 180° space, which might support later reporting of grating orientation and motion direction alike. However, the findings could not explain how a line-like representation (which is invariant to a 180° rotation) would differentiate between opposite motion directions (e.g., up vs down), which was essential for the task at hand. Moreover, if WM representations are abstract and generalised in early sensory areas, where and how would concrete sensory-perceptual detail be retained in the brain, if at all?

An underexplored possibility is that WM representations at different levels of abstraction may coexist within a given area. For instance, EVC can retain visual WM information while concurrently processing perceptual input^9^, suggesting a capacity to multiplex representations with little interference between them (see also^40^, for a similar finding in auditory cortex). Within areas, WM representations were found to undergo substantial transformations over time, indicating that local neural populations can encode retrospective stimulus information and prospective, action-oriented transformations alike^26,41,42^. Individual areas may further hold concurrent WM representations in orthogonal neural formats shaped by priority^43–46^, extending the idea that single neurons can encode a multitude of stimulus- and task variables simultaneously (“mixed selectivity”^47,48^). This raises the possibility that individual areas may hold multilayered WM representations of both abstract and concrete stimulus information. However, many previous studies of visual WM have examined neural representation of a single feature, such as orientation or colour^49,50^, without systematically assessing the representations’ level of abstraction or generality.

Here, we used oriented object stimuli to examine the level of visuospatial abstraction of WM information across the visual cortical hierarchy. Specifically, we asked to what extent orientation representations were object-specific or generalised across visually dissimilar objects, the latter reflecting visuospatial abstraction. We found surprisingly little evidence for a posterior-to-anterior gradient of representational formats from concrete to abstract. Instead, concrete (object-specific) and abstract (object-independent) formats appeared to coexist with orthogonal representational geometries in early visual and higher-level areas alike. Across brain areas, the WM information’s homogeneous level of visuospatial abstraction was topographically dissociable from the emergence of categorical biases, which we observed only in the parietal cortex.

## Results

During fMRI scanning, participants (n = 40) were asked to remember oriented real-world objects in a retro-cued WM task (**Figure 1a**). On each trial, two sample objects were successively presented in random orientations (**Figure 1b**), after which an auditory retro-cue ("one" or "two") informed participants which sample they should remember. After a 12-second delay period, participants were asked to identify the cued object (by toggling between three probe options; see *Methods*) and freely rotate it to report its orientation (continuous report). Behaviourally, participants almost always identified the correct stimulus object [mean percentage correct = 98.2%, s.e.m. = 0.29%; compared to 33.3% chance-level: t(39) = 225.37, p < 0.001, d = 35.63] and reported its orientation with a mean absolute error of 15.9° [s.e.m. = 1.04°; compared to 90° chance-level: t(39) = -70.94, p < 0.001, d = 11.22].

**Fig. 1.**
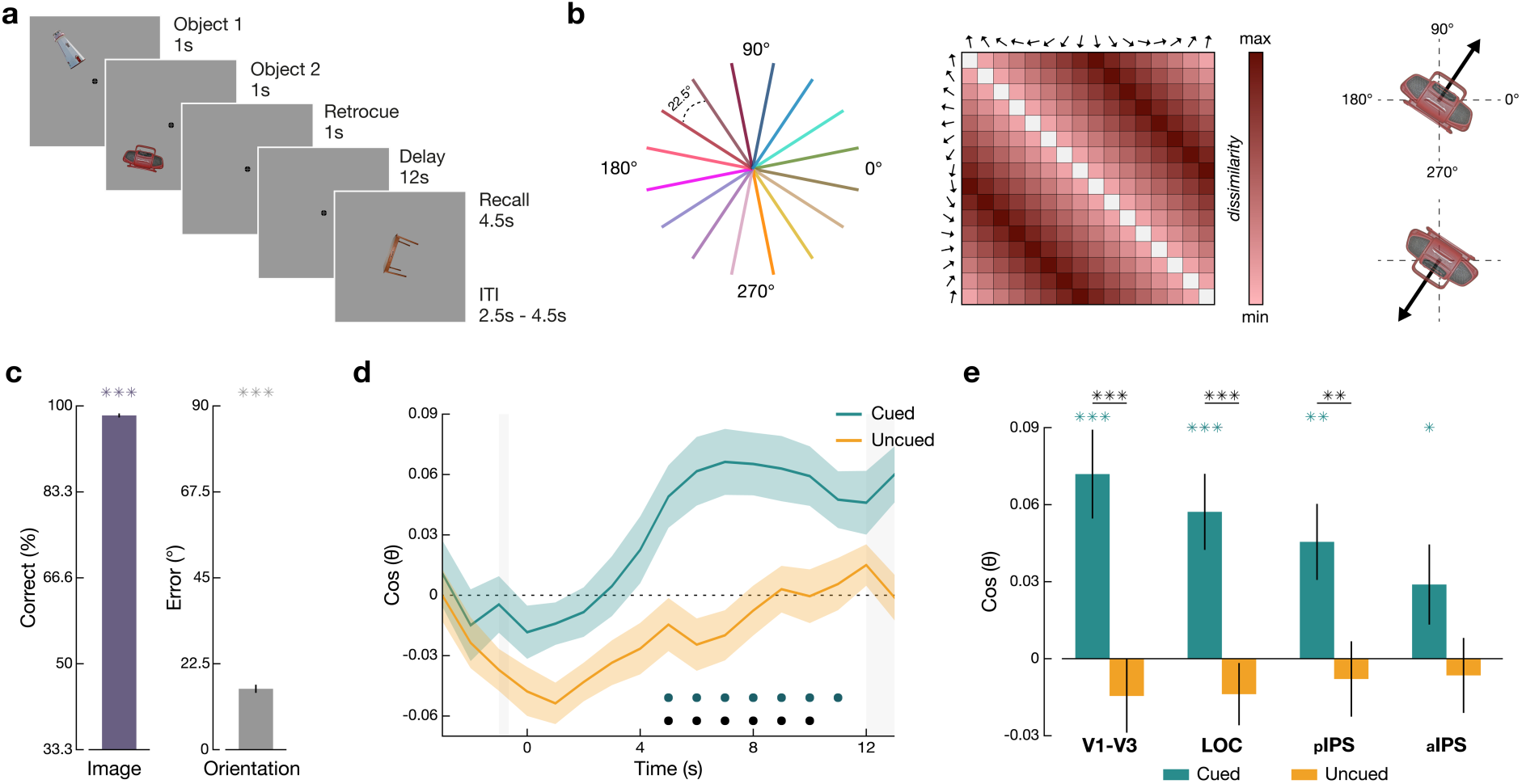
Experimental paradigm and fMRI orientation encoding. **a,** Each trial began with two randomly oriented sample objects presented sequentially at off-centre screen positions, followed by an auditory retro-cue (“one” or “two) indicating which of the two samples was to be remembered. At the test, participants were asked to remember both the identity and orientation of the cued object (see *Methods*). **b,** Left: across trials, each object appeared in 16 evenly spaced orientations. Middle: representational dissimilarity matrix (RDM) of the pairwise angular distances between the 16 possible object orientations (360° model). Right: orientations with opposite directions are maximally distant. **c,** Group behavioural performance. Left: object identification (mean percentage correct), right: orientation report (mean absolute error in degrees). Error bars indicate s.e.m. **d,** Time course of RSA orientation encoding in early visual cortex (V1–3). Vertical shadings indicate the time at which the retro-cue and the recall probe were presented, respectively. During the delay period, the encoding of the to-be-remembered orientation was significantly above zero and significantly stronger than the encoding of the to-be-forgotten orientation. Error shadings show s.e.m., and dots at the bottom indicate time points of significant orientation encoding of the cued object orientation compared to zero (coloured) and compared to the uncued object orientation (black) (p < 0.05, FDR corrected). **e,** Orientation encoding during the delay (4-12 seconds after the retro-cue) in the different ROIs (see also Supplementary Figure S1). Significant encoding of the cued orientation was evident in all ROIs (see Methods). Error bars show s.e.m.; coloured asterisks indicate significance compared to zero, and black asterisks indicate the difference between cued and uncued samples (*p < 0.05, **p < 0.01, ***p < 0.001, FDR corrected), n = 40 participants. LOC: Lateral Occipital Complex; pIPS: posterior intraparietal sulcus; aIPS: anterior intraparietal sulcus.

### Orientation Encoding

We analysed fMRI data from regions of interest (ROIs) that were found to be involved in WM processing of visual orientation information in previous work (V1-V3^7,8,10,39^, LOC^17,37,51–53^, pIPS^8,9,37,39,54^, and aIPS^13,37,54,55^). To examine orientation encoding in these regions, we used an approach based on Representational Similarity Analysis (RSA). Specifically, we compared the pairwise dissimilarity structure of neural activity patterns associated with the 16 sample orientations to that predicted by the angular distances between the samples’ physical orientations (Fig. 1b). We will later refer to this basic orientation model as our “360°” model. The analysis was performed separately for each time point in the WM delay, yielding a time course of orientation encoding (Fig. 1d). For statistical analysis, we focused on the last 8 seconds of the WM delay (i.e., the average dissimilarity pattern between 4 and 12 s after the retro-cue), consistent with previous fMRI studies of WM^8,17,37^.

In the early visual cortex (V1-V3), we observed significant encoding of the cued sample orientation [t(39) = 4.14, p < 0.001, d = 0.66], starting approximately 5s after presentation of the retro-cue (Fig. 1d). The encoding of the cued orientation was significantly stronger than that of the uncued sample [t(39) = 4.55, p < 0.001, d = 0.86, two-tailed], which itself was not significantly above zero [t(39) = -1.02, p = 0.863, d = 0.16]. A qualitatively similar pattern was evident in all ROIs (Fig. 1e and Fig. S1), even though the encoding strength in higher cortical areas (e.g., IPS) appeared somewhat weaker. These results complement previous findings of cue-dependent WM representations in early visual areas^7,10^ and support the idea that WM storage is distributed across the visual cortical hierarchy^9,16,55,56^. An analysis of complementary eye-tracking data recorded in a separate experiment outside the scanner (n = 37) showed that orientation encoding in gaze patterns (cf. Linde ^57^ was minimal with our present task setup (see Supplementary Fig. S5 for details), rendering it unlikely that the robust fMRI encoding was only driven by eye movements (see Discussion).

We then turned to our main analysis question, namely the extent to which the different ROIs encoded the cued stimulus orientation in an object-specific (‘concrete’) or object-independent (generalised or ‘abstract’) format. The stimulus objects used in our experiment differed strongly in visual appearance, but their orientation could be easily transformed into shared rotational coordinates (relative to the object’s real-world upright position; **Figure 2a**, *left*). This allowed us to infer the visual WM representations’ level of abstraction by comparing the orientation encoding within vs between the different objects (**Figure 2a**, *right*). Stronger within- than between-object encoding would indicate a more concrete, object-specific representation, while the relative strength of between-objects encoding reflects the extent to which the orientation information was object-independent (i.e., ‘abstracted’ from the physical stimulus). Note that in our behavioural paradigm, either type of WM representation – concrete or abstract – could in principle be sufficient to successfully perform the task.

**Fig. 2.**
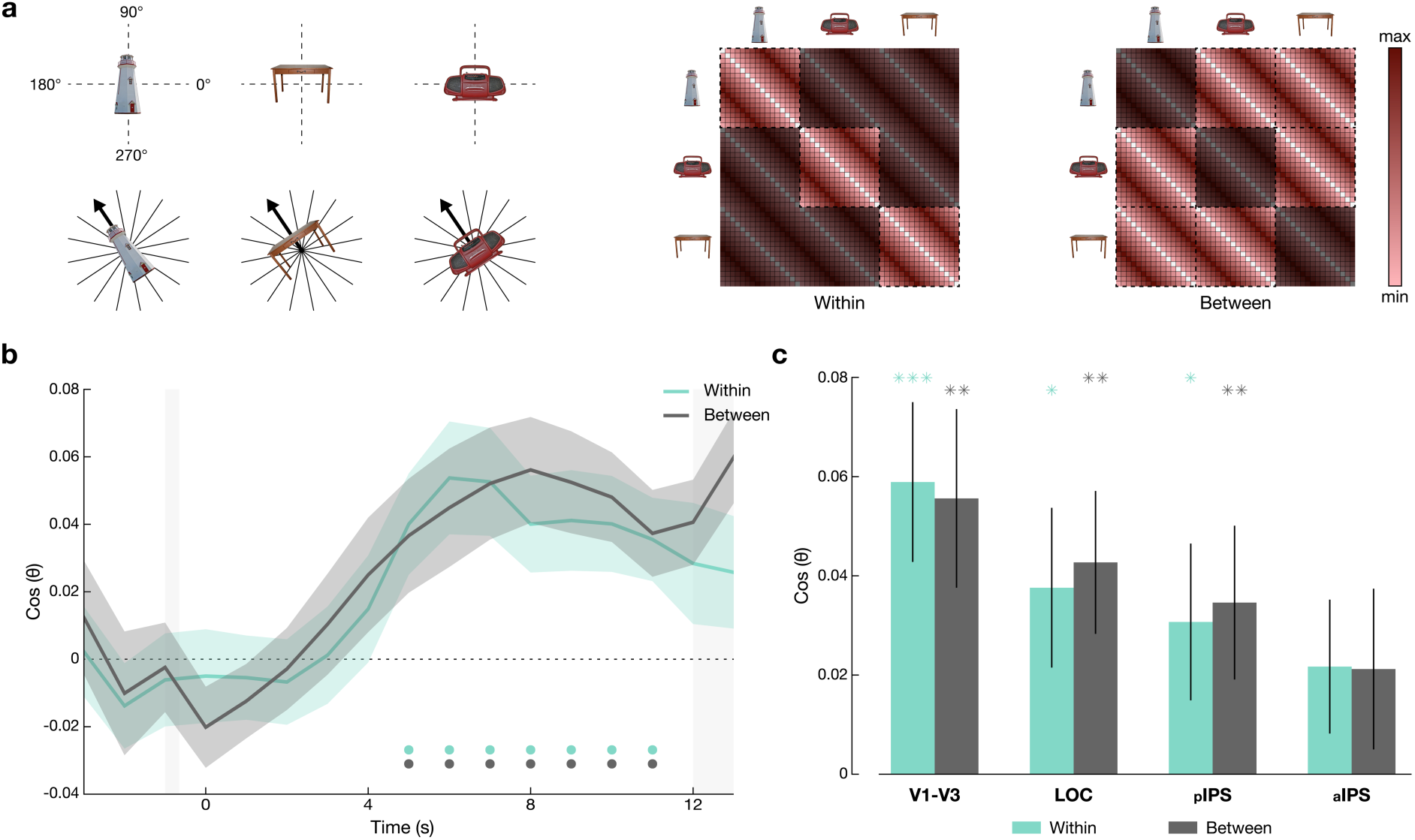
Object-independent orientation encoding. **a**, Left: Examples of stimuli used in the experiment. The stimulus objects differed strongly in visual appearance, allowing us to test the extent to which neural representations of their memorised orientation were generalised (abstracted from visual input). Middle and right: model RDMs of orientation encoding within and between objects. Darkened cells denote distances excluded from each model. **b**, Orientation encoding within and between objects in early visual cortex (V1–V3). The two time courses were nearly identical, indicating that the underlying orientation representation was object-independent. Coloured and grey dots at the bottom indicate time points of significant orientation encoding (p < 0.05, FDR corrected); otherwise, same conventions as Fig. 1d. **c**, Within- and between-objects orientation encoding in the different ROIs. All ROIs showed significant orientation encoding between objects (except aIPS, but see Supplementary Figure S1b), and no difference vs within-objects, indicating object independence (see Results and Supplementary Table S1). Error bars show s.e.m.; coloured and grey asterisks indicate significance compared to zero (*p < 0.05, **p < 0.01, ***p < 0.001, FDR corrected), n = 40 participants.

As expected, orientation encoding within-objects was significant in V1-V3 [t(39) = 3.66, p < 0.001, d = 0.58]. Interestingly, these early visual areas also showed substantial orientation encoding between objects [t(39) = 3.10, p = 0.005, d = 0.49], which was statistically indistinguishable from within-objects encoding [t(39) = 0.15, p = 0.978, d = 0.03]. A Bayes-factor (BF) analysis yielded moderate evidence that within- and between-objects encoding were equally strong (BF_10_ = 0.17), indicating that early visual cortices represented the orientation information in an abstract (object-independent) format. Similarly high levels of abstraction were evident in all ROIs (**Figure 2c**), [LOC: t(39) = -0.24, p = 0.978, d = 0.05, BF_10_ = 0.17; pIPS: t(39) = -0.17, p = 0.978, d = 0.04, BF_10_ = 0.17; aIPS: t(39) = 0.03, p = 0.978, d = 0.01, BF_10_ = 0.17]. Consistently, we found no significant differences in the level of abstraction across ROIs (within/between x ROI interaction; all F(1,39) < 0.25, all p > 0.5; see Supplementary Table S2).

### Line-like (180°) Orientation Encoding

The results thus far suggest that visual areas including V1-V3 encoded the cued WM orientation in an object-independent (“abstract“) format, which does not retain object-specific (“concrete”) visual markers of orientation such as the relative position of the radio’s carry handle, or of the lighthouse’s entrance door (Fig. 1a). However, another form of concrete visual orientation information is how the objects’ overall aspect ratio (as defined by its coarse shape or silhouette) was oriented in space — where the concrete visual memory of a sample’s orientation would resemble that of an oriented “line” (see also ^8^) or “bar” along the object’s physical main axis. If this were the case, the four cardinal axes of our 360° orientation space (up, down, left, right, see Fig. 1b) would collapse into two (vertical, horizontal, Fig. 3a), resulting in a 180° orientation space. Since 180° and 360° circular spaces are orthogonal to each other, a potential line-like (180°) representation may have gone undetected in our above (360°) analysis. Critically, if a line-like (180°) representation reflects concrete visual information, we would expect it to be aligned differently with different objects (e.g., along the height of the lighthouse, but along the width of the radio, reflecting the individual stimuli’s principal axis). Thus, when replacing our previous (360°) analysis model (Fig. 1b) with a 180° model (Fig. 3a), we again compared orientation encoding within vs between objects to infer the level of concreteness (vs abstraction) of the orientation information.

**Fig. 3.**
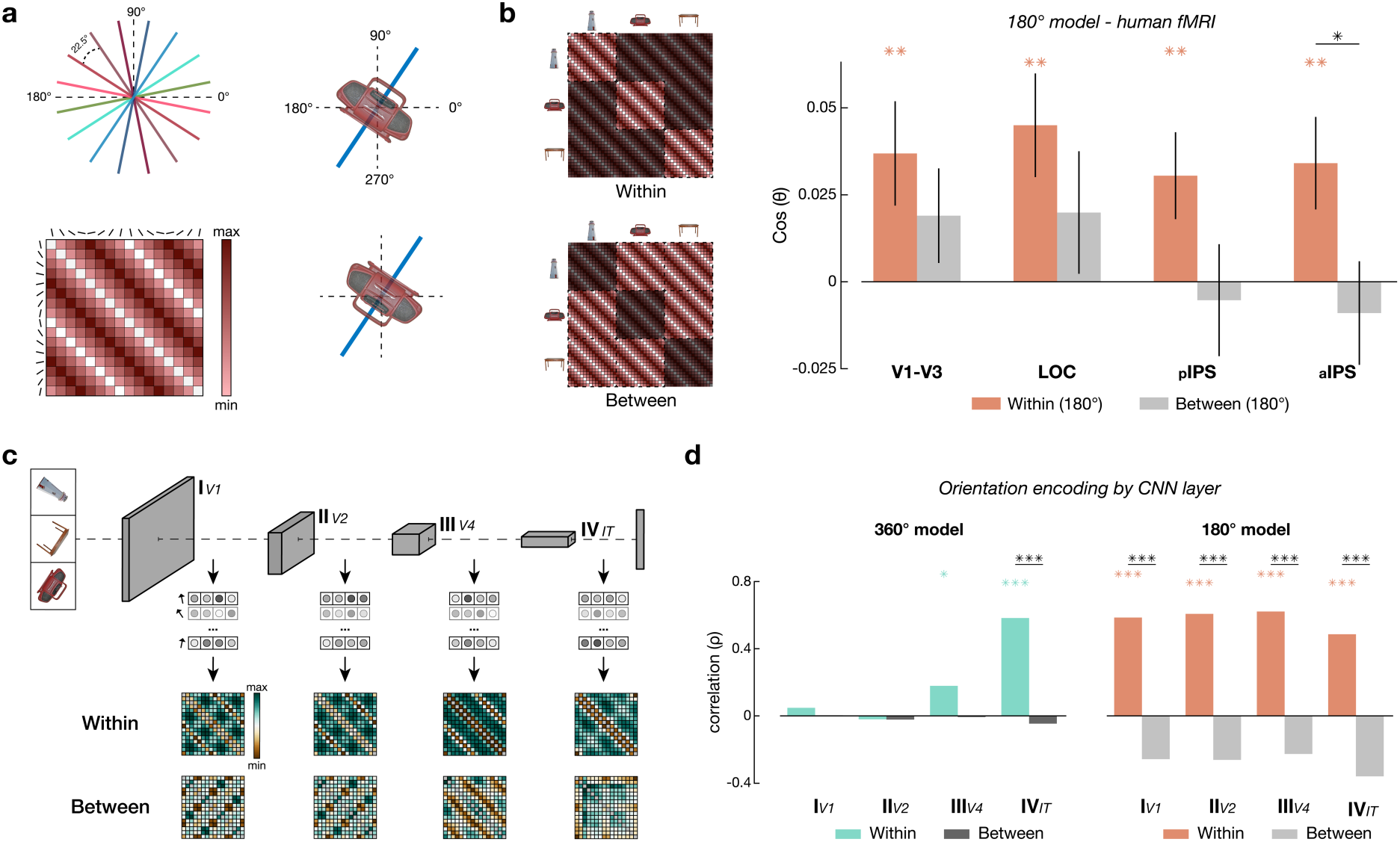
Line-like (180°) orientation encoding and comparison with a deep neural network (CORnet-S). **a,** Illustration of a hypothetical “line-like” (180°) representation of our stimulus orientations (illustrated by coloured lines) and associated model RDM. The model would ignore whether objects are upside-down, but preserve the orientation of their coarse visual shape, or outline. **b,** Neural encoding of 180° orientation during the WM delay. *Left*, model RDMs. *Right,* all ROIs showed significant 180° orientation encoding within objects but not between objects (see Supplementary Table S3 for details). Error bars show s.e.m.; coloured and grey asterisks indicate significance compared to zero (*p < 0.05, **p < 0.01, ***p < 0.001, FDR corrected), n = 40 participants. **c,** CNN analysis approach. We performed RSA analogous to the fMRI analysis on layer activations of a CNN (CORnet-S; see Methods). *Top*, CORnet-S architecture. *Bottom*, layer-specific orientation RDMs within and between objects. **d,** Orientation encoding results for each CNN layer. *Left*: 360° orientation encoding was not generalised (between-objects, *grey*) in any layer, in contrast to the RSA results in fMRI (cf. Fig. 2c). *Right*: 180° orientation encoding within-objects was evident in all model layers. Between-objects encoding was expectedly negative, indicating that the DNN encoded the orientation of the individual objects’ visual main axis (which was perpendicular between, e.g., the radio and the lighthouse, see *top left* in *c*; see *Results*).

Repeating the analysis with a 180° (“line”) version of our orientation model (Fig. 3a) revealed significant within-object encoding of the cued orientation in all ROIs [e.g., V1-V3: t(39) = 2.46, p = 0.009, d = 0.39], whereas between-object encoding was insignificant [V1-V3: t(39) = 1.40, p = 0.265, d = 0.22; see Fig. 3b and Supplementary Table S3 for the remaining ROIs]. While this pattern was evident in all ROIs (Fig. 3b), direct comparison of within- vs between-objects encoding reached statistical significance only in the aIPS [t(39) = 2.32, p = 0.046, d = 0.48; see Supplementary Table S2 for details]. Importantly, the within-object encoding disclosed by our 180° model was also cue-dependent: none of the ROIs showed such encoding for the uncued orientation (all p > 0.05, corrected), and the direct comparison of cued vs uncued orientations was significant in early visual cortex [t(39) = 3.09, p = 0.015, d = 0.63] and LOC [t(39) = 2.54, p = 0.030, d = 0.59]. These results suggest the co-existence of a ‘concrete’ (object-specific) visual WM representation of the physical sample stimuli (in terms of their oriented outline or shape) alongside the ‘abstract’ 360° representation identified in the previous section – intriguingly, within the same set of brain areas.

### Orientation Encoding in an Artificial Neural Network Model

What kind(s) of orientation representation of our stimuli could have been expected to arise in a bottom-up fashion, independent of the specific requirements of our WM tasks? To explore this, we examined the extent to which 360° and/or 180° representations of our stimulus materials emerge in CORnet-S, a hierarchical convolutional neural network (CNN) model of visual object recognition^58^. For each CNN layer, we extracted the activity vectors associated with the different object orientations and compared their representational dissimilarity structure to the predictions of our 360° and 180° models within and between objects (see Methods). Unlike in the human fMRI data (cf. Fig. 2b-c), we found no evidence for generalised (between-objects) 360° orientation encoding in any of the four CNN output layers (Fig. 3d, *left*). Additionally, within-object orientation encoding was only evident in the last layer (IV; mean = 0.608, p < 0.001, see Supplementary Table S4). In contrast, repeating the analysis with our 180° (“line”) orientation model showed robust orientation encoding in all four CNN layers (I-IV; Fig. 3d, *right*). Specifically, the 180°-orientation encoding was strongly positive within objects, but *negative* between objects, as expected if the CNN encoded the orientation of the stimuli’s main visual axis, which systematically differed between objects (see *Methods* and Fig. S3b). In this respect, the CNN patterns resembled our findings with the 180° model in the human fMRI data (cf. Fig. 3b), which likewise encoded the stimuli’s orientation in an object-dependent 180° space.

An important aspect of these observations is that they show 360°- and 180°-orientation encodings to be dissociable in principle, both within and across the different layers of a hierarchical neural network. Furthermore, the CNN findings render it unlikely that the generalised, object-independent (360°) orientation representation we observed in early visual areas (Fig. 2b) was driven in a bottom-up fashion by properties of our specific stimulus materials. While the neural network model did extract object-specific 180° orientation information similar to human visual areas, it failed to produce the generalised 360° orientation information we found multiplexed in those same visual areas during human WM processing.

### Categorial Orientation Biases

Previous studies of visual WM have shown that participants tend to systematically distort memorised orientations away from the cardinal axes (vertical and horizontal), a phenomenon known as ‘cardinal bias’. Repulsive cardinal bias is commonly observed with artificial grating or line stimuli ^17,59^, but has recently been shown to also occur with real-world objects oriented in 360° space ^57^. In further analyses, we asked whether and how such a bias manifested in the geometry of neural activity patterns during the delay period.

We first verified that cardinal repulsion bias was evident in the behavioural data. The participants’ mean signed response errors indeed showed a typical pattern of cardinal repulsion (Figure 4a), in which the orientations were remembered as further away from the cardinal axes than they actually were. Fitting a parameterised geometric model (Linde-Domingo & Spitzer, 2024, see *Methods*), we observed significantly positive values of bias parameter *b* [mean ± s.e.m. = 0.363 ± 0.017, t(39) = 20.93, p < 0.001, d = 3.31], indicating a significant repulsion bias. We then examined the degree to which such a bias also appeared in the delay-period neural representations of 360° orientation in our ROIs. Fitting the bias model to the neural representational geometries (via the associated RDMs, see *Methods*) showed a significantly positive estimate of *b* in aIPS [Figure 4c; mean ± s.e.m. = 0.40 ± 0.13, t(39) = 3.15, p = 0.008, d = 0.50], the most anterior of our ROI. In contrast, *b* was near-zero in the posterior ROIs [V1-V3: mean ± s.e.m. = 0.01 ± 0.13; LOC: mean ± s.e.m. = 0.03 ± 0.13; pIPS: mean ± s.e.m. = 0.09 ± 0.14, *all* t(39) < 1; *all* p > 0.5]. These results replicate and extend recent fMRI findings with oriented grating stimuli ^17^, corroborating that cardinal repulsion bias emerges only at higher levels of the cortical hierarchy. For completeness, we also performed bias analysis with a 180° version (cf. Fig. 3) of the model, which showed no significant effects in any of the ROIs [Figure 4d; *all |b|* < 0.15*; all |*t(39)| < 1.15; *all* p > 0.5].

**Fig. 4.**
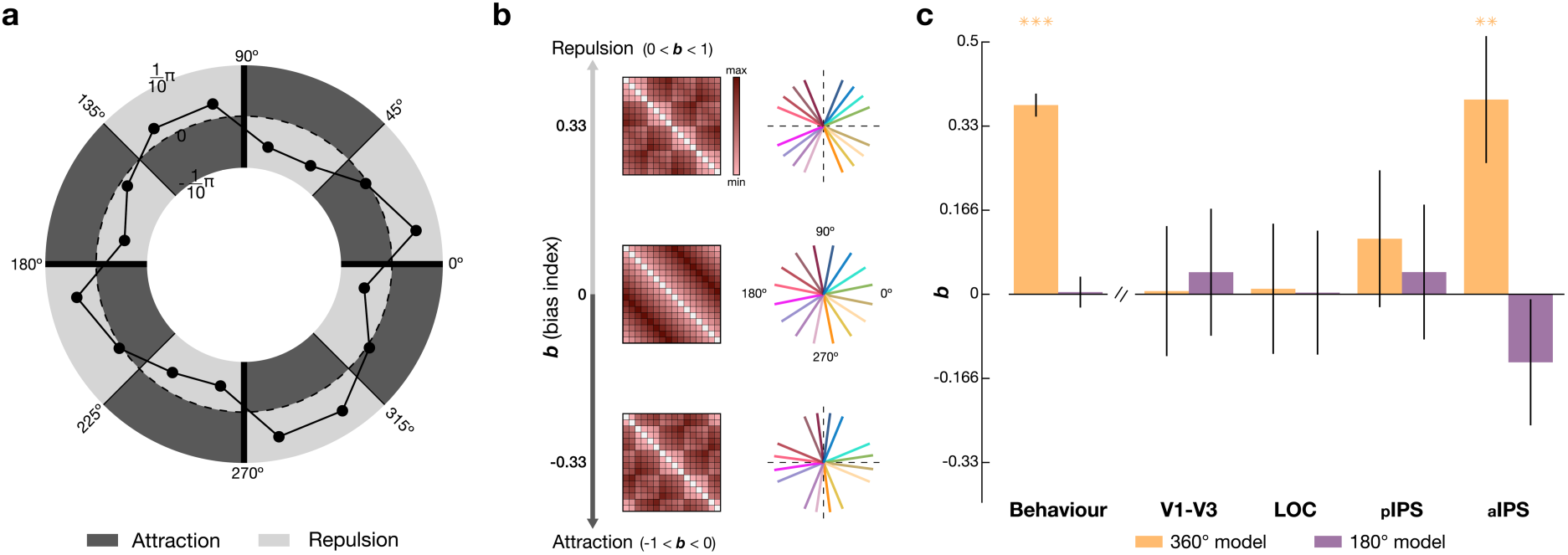
Cardinal repulsion bias. **a,** Behavioural response bias (mean signed response error in radians) for each of the cued target orientations. Dark and light areas indicate whether responses were biased towards (attraction, *dark*) or away from (repulsion, *light*) the nearest cardinal axis. The data showed a clear pattern of repulsion. **b,** We parameterised our orientation models to reflect the level of attractive (*b* < 0) or repulsive bias (*b* > 0) in the model-predicted RDM (see *Methods* and *Results*). **c,** Gold: estimates of *b* from fitting the parameterised 360° model to the behavioural and neural data. A significant repulsion bias was evident in behaviour and in the aIPS. Purple: estimates of *b* from the 180° model showed no significant bias (see Supplementary Fig. S3). Error bars show s.e.m.; coloured asterisks indicate significant differences from zero (*p < 0.05, **p < 0.01, ***p < 0.001, two-tailed, FDR corrected), n = 40 participants.

### Encoding of object identity and spatial location

While the primary focus of our study was on the WM encoding of orientation, we also explored whether the ROIs retained information about the sample object’s identity (e.g., lighthouse, table, or radio) and/or their spatial location on screen (Fig. 1a), independent of orientation. Using multiclass decoding (see *Methods*) we found information about the cued object’s identity in all four ROIs [Supplementary Fig. S4a and Table S7; e.g., V1-V3: t(39) = 3.68, p < 0.001], whereas the classification accuracy for the uncued object was not significantly above chance level [e.g., V1-V3: t(39) = -0.59, p = 0.758]. Direct comparison between the cued and uncued objects showed significant differences in all regions except the aIPS [V1-V3: t(39) = 3.11, p = 0.005; see Supplementary Fig. S4a and Table S7 for the remaining ROIs]. Thus, in addition to multilayered orientation information, these regions also held information about the to-be-reported stimulus object^27^.

Lastly, again using RSA, we found that the spatial location of the cued sample (Fig. 1a, *left*) was encoded during the WM delay in all ROIs except aIPS [Supplementary Figure S4b and Table S8; e.g., V1-V3: t(39) = 3.73, p < 0.001, d = 0.59], and its encoding was significantly stronger than that of the uncued sample in V1-V3 [t(39) = 2.55, p = 0.03, d = 0.53] and LOC [t(39) = 3.40, p = 0.008, d = 0.70; see Supplementary Fig. S4b and Table S8 for the remaining ROIs]. Thus, these early areas also retained information about the cued stimulus’s spatial location on screen even though participants were never asked to report it, extending previous findings in other WM tasks^60–62^.

## Discussion

To summarise our main findings, representational geometry analysis during cued WM maintenance showed evidence for visuospatial abstraction of orientation information not only in parietal and extrastriate areas, but also in early visual cortex (V1-V3). In addition, within the very same areas, an orthogonal, line-like representation of the WM sample’s concrete visual shape (or outline) was evident. Together, the results suggest the widespread coexistence of abstract and concrete visual WM information in distributed WM storage, with surprisingly little variation between areas.

Since the seminal discovery of visual WM information in primary visual cortex^7,10^, the precise nature and role of mnemonic representation in early sensory areas have remained subject to debate^16,63–66^. While it has been implied that early sensory areas may retain concrete sensory detail, e.g., in terms of sustained perceptual patterns^18^ or through top-down driven reconstruction of a high-dimensional representation^67^, evidence has emerged that visual WM information in early visual cortex can be more abstract and low-dimensional than previously thought^8,25,29,39^. Our finding of object-independent orientation information in V1-V3 supports the latter view. Rather than a pictorial representation of the just-seen visual image, V1-V3 appeared to maintain a visuospatial abstraction, potentially reflecting a generic direction for turning the probe object at test. This high level of visuospatial abstraction was evident even though our task required remembering both the cued object’s orientation and its identity, which might have been expected to promote a concrete-visual memory of the rotated sample image.

The generalised encoding of object orientation in V1-V3 coexisted with an orthogonal, ‘line-like’ representation of the sample image’s orientation in a 180°-rotational space. With our selection of object stimuli, the 180° encoding may be interpreted as a blurred visual representation of the concrete image’s oriented outline or shape, which does not retain perceptual or semantic information that would distinguish between the object’s top or bottom. The coexistence of the two orthogonal formats (360° and 180°) was not limited to V1-V3 but was topographically widespread, including in LOC and IPS. Importantly, unlike the object-independent 360° encoding, the 180° encoding showed little generalisation across objects, consistent with an interpretation as concrete-visual information about the sample image. Together, the results suggest that during cued WM maintenance, areas from early visual to parietal cortex retained both a goal-oriented visuospatial abstraction of the objects’ “semantic” orientation (relative to their real-world upright position) alongside coarse information about the sample image’s concrete visual appearance, with surprisingly little variation across the visual cortical hierarchy. A potential explanation for these findings is multiplexing, where different aspects are encoded simultaneously within separable subspaces of neural activity^43^. Multiplexing could be maintained by different neuronal subpopulations or by overlapping populations that encode multiple features in orthogonal activity patterns^47,48^. Alternatively, or additionally, the different aspects of a multilayered WM representation might be multiplexed in time, by intermittent or rhythmically alternating population codes^68–70^. While such mechanisms have been primarily characterised in the prefrontal cortex^47,48,71^, they may extend to sensory regions, including the early visual cortex^9,72^.

Examining our oriented object stimuli with an artificial neural network model of visual processing (CORnet-S), we observed encoding of object orientation in a 180° space. This encoding was object-specific and distributed across all network layers, similar to our human fMRI finding of 180°-orientation information during WM processing. However, in clear distinction from the fMRI results, the CNN showed no indication of abstract-generalised orientation encoding in 360° space. While CORnet-S was not trained on orientation processing^58^, we derive two implications from these complementary modelling results: (i) encoding our stimuli’s orientations in an object-specific 180° space appears as a plausible visual coding scheme, and (ii) 180° and 360° encoding of our stimuli are dissociable in principle (i.e. one can occur without the other) in a hierarchical neural network. At the same time, the findings corroborate that during WM processing, human visual areas can host information in abstract-generalised formats that do not trivially emerge from a basic (mostly feed-forward) architecture of visual processing.

While the orientation representations’ level(s) of generalisation, as well as the coexistence of object-specific and object-independent formats, were remarkably homogenous across our ROIs, we observed regional differences in the extent to which the representations were categorically biased. Recent work using visual gratings or colour patches showed that categorical bias is absent in EVC but emerges in extrastriate and parietal areas^17,30^, as well as in the precentral sulcus^29^. Broadly consistent with this, we found categorical bias (cardinal repulsion) only in the most anterior of our ROIs, the aIPS. The relatively late processing stage might reflect that with our real-world object stimuli, cardinal orientation bias in 360° space might be driven by relatively high-level concepts^57,73,74^, such as upright vs upside down, which rely on a semantic notion of the object and its natural upright position ^57^. Interestingly, we found no evidence for bias in the concurrent ‘line-like’ (180°) orientation representation, potentially indicating that categorical distortions only affect aspects that are directly relevant for the task at hand (here, orientation in 360° space; cf. findings in 180° orientation tasks ^17,29,30^).

It is increasingly recognised that visuospatial WM processing can be accompanied by miniature eye movements^57,75–77^. In particular, with central stimulus presentation at fixation, the to-be-maintained orientation of a WM sample was found to induce small but systematic gaze deflections throughout sustained WM delays^57,76^. Our complementary eye-tracking results showed that with off-centre sample presentation at varying screen locations (Fig. 1a, *left*), orientation encoding in gaze was largely absent (see Supplementary Fig. S5 for details), rendering it unlikely that our robust fMRI findings could be explained solely by systematic eye movements. Consistently, supplementary fMRI analysis showed no 360° or 180° orientation encoding in the Frontal Eye Fields (FEF; Supplementary Fig. S6), which would be associated with spatial attention and saccade planning^78^. A cautionary note remains that even with off-centre presentation, we could still detect residual orientation encoding in our dedicated eye-tracking recordings outside the scanner, but only when aggregating a manifold more task trials than were recorded in the fMRI experiment (see Supplementary Fig. S5 for details). As such, orientation encoding in gaze appeared too weak to be considered a primary explanation for our fMRI results.

In addition to the multiplexing of object-independent and object-specific orientation information across the visual cortical hierarchy, the same areas also retained information about the cued object’s identity (e.g., lighthouse vs table vs radio), which was also to be reported at the WM test. While this additional finding adds to the notion of multilayered WM representation within areas, we prefer not to interpret it in terms of levels of abstraction; the object’s identities could have been distinguished by low-level perceptual (e.g., colour) or semantic features (e.g., building vs furniture) or both. Finally, consistent with prior EEG work^60,79,80^, we found sustained encoding of the cued sample’s physical location on screen (Supplementary Fig. S4b), even though the location was task-irrelevant (i.e., not to be reported at the WM test). Interestingly, unlike the task-relevant WM information, sample location was only reflected in early ROIs (V1-V3 and LOC), which may suggest that the task-irrelevant information was not fed into higher-order (e.g., parietal) decision circuits.

Why would distributed WM maintain such multilayered representation(s) of concrete and abstract formats alongside task-irrelevant information? WM has been characterised as the goal-directed transformation of retrospective stimulus information into a task-appropriate prospective response^16,81^. Sustained WM representations in visual cortices are thought to rely on top-down feedback from fronto-parietal regions^13,18,28^ and to involve formats distinct from those used in perception^9,39,50^. The widespread coexistence of concrete-retrospective and abstract, task-oriented formats may support WM’s flexibility in providing memoranda at the level of abstraction that best fits the current task^67^, even when task requirements inadvertently change^82^. Future studies using recordings with higher temporal precision may help understand the fine-grained dynamics underlying such representational flexibility.

## Methods

### Participants

We recruited n = 46 participants (42 right-hand dominant, 24 female, 22 male, mean age: 25.3 ± 4.8 years) through the Max Planck Institute for Human Development’s participant database. We excluded three participants who did not complete the experiment and an additional three due to excessive head movement, leaving n = 40 participants for analysis (37 right-hand dominant, 21 female, 19 male, mean age: 25.7 ± 4.6 years). All participants met the following criteria: normal or corrected-to-normal vision, age 18-35 years, fluency in English or German, no known psychiatric disorders or use of centrally active drugs, and no prior knowledge of the research questions. The study was approved by the ethics committee of the German Psychological Society (DGPs). Participants provided written informed consent and were reimbursed with €12 per hour, with an additional €5 bonus upon completion. The experiment used German or English for visual and audio instructions, depending on the participant’s preference (35 chose German).

### Stimuli

We selected nine colour images of everyday objects from the BOSS (Birmingham Object Recognition Battery) database ^83^. Specifically, the selection included three objects taller than wide, three wider than tall, and three with approximately equal height and width. We then divided these into three sets, each containing three objects with distinct aspect ratios (Fig. S3a). We assigned one set to each participant, balancing set frequency across the group. The images were cropped and resized to approximately 3° visual angle, based on in-scanner screen measurements of the distance from the screen to the participant’s eyes (approx. 96.5 cm).

### Instructions and training

After providing informed consent and before starting the scan, participants received a briefing about magnetic resonance imaging and an introduction to the experimental tasks. They then had a short training inside the MRI scanner without image acquisition, which included five instruction trials (with shorter retention intervals and response feedback) and five practice trials with the same specifications as the experiment’s trials (without feedback).

### Task and Procedure

The experiment comprised eight functional runs, each containing 24 trials (192 trials in total). Each trial began with a fixation dot (0.6° diameter) displayed centrally for 1000 ms. Two sample objects appeared sequentially, each in a random orientation drawn from 16 equidistant angles (11.25° to 348.75°), which excluded the cardinal axes (0°, 90°, 180°, 270°, see Fig. 1b). To minimise orientation-dependent eye movements in the scanner, we presented the sample stimuli at four off-centre locations, each 3° distant from the fixation point (Supplementary Fig. S3b). Throughout the experiment, each sample object appeared four times in each orientation, once in each location, resulting in a full counter-balancing of the screen locations.

Each object appeared for 500 ms, followed by a 500 ms blank screen. The two objects on each trial were chosen pseudorandomly without replacement. An auditory retro-cue ("one" or "two") indicated which object participants should remember, followed by a 12-second delay period. After the delay, a probe object (randomly drawn from the participant’s stimulus set) appeared at the centre of the screen in a random orientation. Participants were asked to select the correct object by toggling between the three possible objects, then rotated the selected object to match its memorised orientation using MR-compatible response buttons. Trials in which participants failed to respond within 4500 ms were excluded from the behavioural analysis (0.42% of trials on average). After participants submitted their response, a variable inter-trial interval ensued (2500-4500 ms, uniformly randomly varied). We instructed participants to maintain their gaze on a centrally presented fixation dot throughout the trial periods. The experiment was run using PsychoPy-3 in MATLAB 2017a (MathWorks). The visual stimuli were displayed in-scanner on a 48x34 cm screen (1920x1080 px resolution, 60 Hz frame rate), and the auditory cues were played through OptoACTIVE Earphones (Optoacoustics).

### Neuroimaging data acquisition

We acquired anatomical and functional brain imaging data using a 3T Siemens Magnetom Tim Trio MRI scanner with a 32-channel Head Matrix Coil at the Max Planck Institute for Human Development, Berlin, Germany.

#### Functional data

We collected functional scans using a multi-band (MB) imaging protocol (factor of 3) with the following parameters: TR = 1000ms, TE = 27ms, FA = 61°, 66 × 66 imaging matrix, field of view (FOV) = 192 mm^2^, in-plane resolution = 3 mm^2^. We acquired 45 axial slices (3 mm thickness, 0 mm gap) for full cortex coverage. Each run included 567 functional volumes, with a total run time of 9 minutes and 34 seconds. We also collected two field map scans (Spin Echo images with anterior-posterior and posterior-anterior phase-encoding directions) to measure signal homogeneity.

#### Anatomical data

We collected High-resolution T1-weighted anatomical images using an MP-RAGE sequence with the following parameters: FOV = 256 mm^2^, 192 axial slices, with an in-plane resolution of 1 mm^2^, 9° flip angle (FA), and 900 msec Inversion time (TI).

### Preprocessing

Preprocessing was performed using *fMRIPrep* 23.0.2^84^, which is based on *Nipype* 1.8.6^85,86^.

#### Preprocessing of B_0_ inhomogeneity mappings

A *B_0_*-nonuniformity map (or *fieldmap*) was estimated based on two echo-planar imaging (EPI) references with topup^87^.

#### Anatomical data preprocessing

The T1-weighted (T1w) image was corrected for intensity non-uniformity using ANTs 2.3.3^88,89^ and was used as a T1w-reference throughout the workflow. The T1w reference was skull-stripped with a Nipype implementation of ANT. Brain tissue segmentation of cerebrospinal fluid (CSF), white matter (WM), and grey matter (GM) was performed on the brain-extracted T1w^90^. Volume-based spatial normalisation to one standard space (MNI152NLin2009cAsym) was performed through nonlinear registration using brain-extracted versions of both the T1w reference and the T1w template. The ICBM 152 Nonlinear Asymmetrical template version 2009c template was used for spatial normalisation and accessed with TemplateFlow 23.0.0^91^.

#### Functional data preprocessing

First, a reference volume and its skull-stripped version were generated using a custom methodology of fMRIPrep. Head-motion parameters concerning the blood-oxygen-level-dependent (BOLD) reference are estimated before any spatiotemporal filtering using FSL^92^. The estimated fieldmap was then aligned with rigid registration to the target EPI (echo-planar imaging) reference run. The field coefficients were mapped onto the reference EPI using the transform. BOLD runs were slice-time corrected using AFNI^93^. The BOLD reference was then co-registered to the T1w reference using bbregister (FreeSurfer), which implements boundary-based registration^94^. Co-registration was configured with six degrees of freedom. Confounds for framewise displacement and three region-wise global signals time-series were calculated on the preprocessed BOLD. Framewise displacement was computed for each functional run using two formulations^92,95^, and the three global signals were extracted within the CSF, the WM, and the whole-brain masks. The confounds’ time series derived from head motion estimates and global signals were expanded with the inclusion of temporal derivatives and quadratic terms for each^96^. The BOLD time series were resampled into standard space, generating a preprocessed BOLD run in MNI152NLin2009cAsym space. All resamplings were performed with a single interpolation step by composing all the pertinent transformations. In contrast, Gridded (volumetric) resamplings used Lanczos interpolation to minimise the smoothing effects of other kernels^97^.

#### Head-Movement exclusion criteria

We excluded participants with excessive head movement based on the following thresholds: maximum translation/rotation exceeding 3°/3 mm, mean translation/rotation exceeding 1.5°/1.5 mm, and mean framewise displacement exceeding 0.25°/0.25 mm.

### Regions of Interest (ROI) selection

We used the Julich-Brain probabilistic cytoarchitectural map^98^, a 3d atlas of the human brain’s cytoarchitecture, to precisely identify areas commonly associated with WM maintenance in humans^7,8,13,17,54^. Based on this atlas, we parcellated four regions of interest: Early visual cortex (V1-V3); Lateral Occipital Complex (LOC); posterior intraparietal sulcus (pIPS); anterior intraparietal sulcus (aIPS) (see supplementary table for area names and their corresponding Julich atlas sub-regions). For our General Linear Model (GLM) analysis, we selected voxels from each ROI by combining the Julich atlas with individual probabilistic grey matter masks generated by the fMRIPrep preprocessing pipeline. First, we masked the functional images (in MNI space) using the ROI parcellation from the Julich atlas, with a threshold of 0.1. For each participant, we masked the resulting image with their corresponding grey-matter mask calculated during preprocessing (using a similar threshold).

### fMRI GLM specification

The preprocessed functional data were initially z-transformed (signals and confounds) on a per-voxel and per-run basis. To obtain the neural response time series for each trial, we estimated the BOLD response at every time point from the first stimulus’s onset to the end of the probe period (19 time points in total). The univariate estimation was performed using GLM with a Finite Impulse Response (FIR), which models the time series with a set of time-shifted regressors. The FIR method is particularly well-suited for examining fMRI signals during delay periods as it does not impose constraints on the shape of the hemodynamic response. Additionally, we included confound regressors for motion (six regressors), CSF, WM, global signals, their temporal derivatives, quadratic terms, and squares of derivatives^99^, as well as six discrete cosine-basis regressors for low-frequency physiological and scanner noise. The GLM was estimated separately for each functional run, and residuals were recovered for subsequent noise normalisation. We performed no additional standardisation, signal scaling, or detrending of the data. The GLM analysis was carried out using a combination of custom-made functions and the built-in pipeline in Nilearn 0.11.1^100^.

### Multivariate Representational Similarity Analysis (RSA)

RSA of the fMRI data was performed separately for each participant using cross-validation^101^. First, we applied univariate noise normalisation to the data using the standard deviation of the GLM residuals^102^. Next, at each time point, we calculated the pairwise Euclidean distances between the activity patterns for each stimulus condition (16 orientations x 3 objects). Specifically, we estimated the squared Euclidean distance *d* between activity patterns (*V*) for stimulus conditions *x* and *y* (*x* ⧣ *y*) over M data partitions as follows:

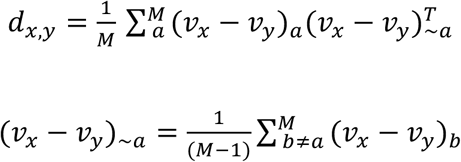

Where the pattern difference from run *a* is multiplied by the pattern difference averaged over all runs except *a* (*∼a*)^103^. Applied to all stimulus conditions and averaged across folds, the procedure yields a 48 x 48 Representational Dissimilarity Matrix (RDM) for each participant at each time point^101^. To obtain a bilateral estimate, we calculated the RDMs for each ROI in the left and right hemispheres separately, then averaged the results for homologous regions^102^. The procedures above were performed separately for the cued and the uncued WM sample (cf. Fig. 1).

To examine orientation encoding during the WM delay, we computed the whitened cosine similarity between the participant-specific neural data RDM and a model RDM^104^. Our basic orientation model was a simple circle geometry, with the model RDM consisting of the pairwise angular distances between 16 evenly spaced angles around the circle (‘360 model’). For analysis of cued vs uncued orientation encoding (Fig. 1), the data RDM (48 x 48, comprising distances within and between objects) was averaged across objects into a 16 x 16 RDM. For subsequent analyses (Fig. 2), we averaged the distances within and between objects to obtain separate RDMs. RSA was performed at each time point, yielding a time course of orientation encoding (e.g. Fig. 1d). For statistical analysis, the data RDMs were first averaged across time points between 4 and 12 seconds after the retro-cue. This time window aligns with previous fMRI studies of visual WM and accounts for delays in hemodynamic responses^8,9,17,37,105^. The RSA analyses were performed in Python using RSAtoolbox 0.2.0^106^.

### Statistics

We used one-tailed t-tests to test whether group means were greater than zero and paired two-tailed t-tests when comparing submodels (e.g., within- vs between-object models). To correct for the number of statistical tests performed across the four ROIs, we used the False Discovery Rate (FDR)^107^.

### Orientation encoding in CNN

When examining orientation encoding in CORnet-S, we used the same object images and orientations that were shown to participants in the fMRI experiment. We followed standard image preprocessing for CORnet-S^58^, resizing images to 256×256 pixels, applying centre cropping to 224×224 pixels, and converting them into tensors. Each image was normalised using ImageNet mean values [0.485, 0.456, 0.406] and standard deviations [0.229, 0.224, 0.225]. We then extracted activations from four output layers (V1, V2, V4, IT) and analysed the activation patterns using RSA. To do so in a manner similar to the fMRI data (where participants were assigned to different stimulus sets), we performed RSA for each of the three stimulus subsets separately and averaged the results. Unlike in the fMRI analysis, we did not perform cross-validation because CORnet-S activations lack measurement variance. For the same reason, we used whitened Pearson correlation (rather than cosine similarity) to quantify the similarity between layer RDMs and model RDMs^101,108^. Model similarity was quantified for each object pairing, then averaged. To assess the statistical significance of the CNN results, we used a percentile test with bootstrapping^109^. For each statistical inference, we created 20,000 random permutations of the model RDM (i.e., with shuffled conditions) and evaluated their Pearson correlation with the data RDM. We then compared the result obtained with the original model RDM against the bootstrapped distribution. The p-value indicates the percentile of bootstrap results that are smaller and/or larger than the correlation between the data RDM and the unpermuted model. We corrected the resulting p-values for multiple comparisons across the four network layers using FDR.

### Bias modelling

To examine whether behavioural responses and neural activity patterns for orientations exhibited a bias towards or away from the cardinal axes, we adopted a modelling approach similar to Linde-Domingo and Spitzer^57^. First, we defined two extreme scenarios where each of the 16 orientations was shifted to the nearest cardinal (0°, 90°, 180°, or 270°; *M*_attract_) or diagonal (45°, 135°, 225°, or 315°; *M*_repulse_). We then parametrically distorted the original orientations (*M*_0_) towards either *M*_attract_ or *M*_repulse_ by weighted summation:

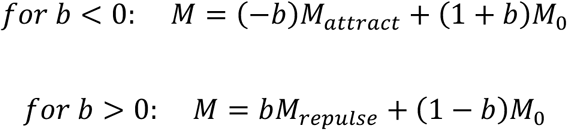

Where the weight parameter *b* (ranging from -1 to 1) reflects the strength of attraction (*b* < 0) or repulsion (*b* > 0). Figure 4b illustrates the model RDMs (pairwise angular distances) resulting from exemplary values of *b*. Note that *b* = 0 equals the base model (*M*_0_, i.e. no bias). To estimate bias in behaviour, the parameterised model RDM was fitted to the behavioural data RDM derived from the participant’s mean orientation reports. For the neural data, the model RDM was fitted to the neural data RDMs in the different ROIs. In both cases, the model was fitted using exhaustive search (-1 to 1, with a step size of 0.01) to identify the value of *b* that maximised the cosine similarity between the model and data RDMs (one behavioural and four brain ROIs; see Fig. 4c).

### Object identity decoding

To decode the sample objects’ identities (e.g., table, lighthouse, or radio, see Supplementary Fig. S3) independent of their orientation, we used a regularised logistic regression classifier implemented in Scikit-learn^110^. Since each participant’s stimulus set contained three possible images, we carried out a three-way multiclass classification using a one-vs-all strategy. Decoding was performed for cued and uncued objects separately using leave-one-run-out cross-validation. At each fold, the decoder was trained on data from seven of the eight runs and tested on the remaining run. The data for each ROI were obtained using the FIR-GLM (see above) and averaged over the period from 4 to 12 seconds after the retrocue. Classification accuracy was evaluated at the group level against a chance level of 33.33% (three possible objects).

### Supplementary eye-tracking data

For technical reasons, no eye-tracking data were recorded during the experiment in the scanner. However, to examine the possibility that fMRI activity patterns in our task might have been confounded by orientation-dependent eye movements^57,111^, we analysed eye-tracking data from a separate WM experiment performed outside the scanner (Supplementary Figure S5). For this experiment, we recruited a separate sample of 40 participants (23 female, 16 male, 1 diverse, mean age = 26.2 ± 3.8 years), of whom n = 3 (1 female and 2 male) were excluded (2 due to incomplete data files and 1 due to insufficient data quality). All participants provided written informed consent, and the experiment was approved by the ethics committee of the German Psychological Society (DGPs). The task design and procedure closely resembled the fMRI experiment, but with a shorter delay period (3600 ms), and each participant performed a substantially larger number of trials (576). Throughout the experiment, monocular gaze position was recorded using a desktop-mounted EyeLink 1000 eye tracker (SR Research) at a 500 Hz sampling rate. Preprocessing of the eye-tracking data closely followed the procedures described in Linde-Domingo & Spitzer (2024), except we used a more lenient artefact threshold (500 px) to account for possible orientation encoding in larger eye movements. For comparability with the fMRI experiment, we subsampled 192 trials from each participant (the first 4 occurrences of each unique combination of orientation and object identity for both the cued and uncued items) to match the trial numbers and stimulus conditions used in fMRI.

## Data availability

The processed fMRI data generated in this study are publicly shared at https://osf.io/nu2ex/overview?view_only=ade1cf4d74c44dbda6b9b8c3a7c89bbd. The data include the beta coefficients (effect sizes) for each voxel within an ROI at each time point in each trial.

## Code availability

The code required to replicate all figures and results from the data is available at https://arc-git.mpib-berlin.mpg.de/yizhar/memfmri_analysis/. The code and stimuli used for the experiment are available at https://arc-git.mpib-berlin.mpg.de/yizhar/memorfmri.

## Author Contributions

Conceptualization: OY, BJS; Data curation: OY, FB, IPS; Formal Analysis: OY, FB, IPS, FBr; Funding acquisition: BJS; Investigation: OY, IPS; Methodology: OY, BJS; Software: OY, FB, IPS, FBr; Supervision: BJS; Validation: OY; Visualization: OY; Writing – original draft: OY, BJS; Writing – review & editing: OY, FB, IPS, FBr, BJS

## Acknowledgements

We thank Sonali Beckmann and Nadine Taube from the MRI unit at MPIB for their assistance in implementing the study. We are grateful to Anouk Bielefeldt, Jiani Ge, Tara Beilner, and Aleksandra Zinoveva for their help with data collection, and to Juan Linde-Domingo for his contribution to the supplementary experiment.

This research was supported by European Research Council Consolidator Grant ERC-2020-COG-101000972 (B.S.) and Deutsche Forschungsgemeinschaft (DFG) Grant 462752742 (B.S.). The funders had no role in the study design, data collection and analysis, or decision to publish.

## References

1. Funahashi, S., Bruce, C. & Goldman-Rakic, P. Dorsolateral prefrontal lesions and oculomotor delayed-response performance: evidence for mnemonic ‘scotomas’. J. Neurosci. 13, 1479–1497 (1993).

2. Miller, E. K., Erickson, C. A. & Desimone, R. Neural Mechanisms of Visual Working Memory in Prefrontal Cortex of the Macaque. J. Neurosci. 16, 5154–5167 (1996).

3. Fuster, J. M. Network memory. Trends in Neurosciences 20, 451–459 (1997).

4. Bisley, J. W., Zaksas, D. & Pasternak, T. Microstimulation of Cortical Area MT Affects Performance on a Visual Working Memory Task. Journal of Neurophysiology 85, 187–196 (2001).

5. Constantinidis, C. & Steinmetz, M. A. Neuronal activity in posterior parietal area 7a during the delay periods of a spatial memory task. Journal of Neurophysiology 76, 1352–1355 (1996).

6. Pasternak, T. & Greenlee, M. W. Working memory in primate sensory systems. Nat Rev Neurosci 6, 97–107 (2005).

7. Harrison, S. A. & Tong, F. Decoding reveals the contents of visual working memory in early visual areas. Nature 458, 632–635 (2009).

8. Kwak, Y. & Curtis, C. E. Unveiling the Abstract Format of Mnemonic Representations. Neuron 1–7 (2022) doi:10.2139/ssrn.3987488.

9. Rademaker, R. L., Chunharas, C. & Serences, J. T. Coexisting representations of sensory and mnemonic information in human visual cortex. Nature Neuroscience 22, 1336–1344 (2019).

10. Serences, J. T., Ester, E. F., Vogel, E. K. & Awh, E. Stimulus-Specific Delay Activity in Human Primary Visual Cortex. Psychol Sci 20, 207–214 (2009).

11. Naughtin, C. K., Mattingley, J. B. & Dux, P. E. Distributed and Overlapping Neural Substrates for Object Individuation and Identification in Visual Short-Term Memory. Cereb. Cortex bhu212 (2014) doi:10.1093/cercor/bhu212.

12. Yu, Q., Panichello, M. F., Cai, Y., Postle, B. R. & Buschman, T. J. Delay-period activity in frontal, parietal, and occipital cortex tracks noise and biases in visual working memory. PLoS Biol 18, e3000854 (2020).

13. Bettencourt, K. C. & Xu, Y. Decoding the content of visual short-term memory under distraction in occipital and parietal areas. Nat Neurosci 19, 150–157 (2016).

14. Christophel, T. B., Hebart, M. N. & Haynes, J.-D. Decoding the Contents of Visual Short-Term Memory from Human Visual and Parietal Cortex. J. Neurosci. 32, 12983–12989 (2012).

15. Ester, E. F., Sprague, T. C. & Serences, J. T. Parietal and Frontal Cortex Encode Stimulus-Specific Mnemonic Representations during Visual Working Memory. Neuron 87, 893–905 (2015).

16. Christophel, T. B., Klink, P. C., Spitzer, B., Roelfsema, P. R. & Haynes, J. D. The Distributed Nature of Working Memory. Trends in Cognitive Sciences 21, 111–124 (2017).

17. Chunharas, C., Wolff, M. J., Hettwer, M. D. & Rademaker, R. L. A gradual transition toward categorical representations along the visual hierarchy during working memory, but not perception. eLife 10.7554/eLife.103347.1 (2025) doi:10.7554/eLife.103347.1.

18. D’Esposito, M. & Postle, B. R. The Cognitive Neuroscience of Working Memory. Annu. Rev. Psychol. 66, 115–142 (2015).

19. Sreenivasan, K. K., Curtis, C. E. & D’Esposito, M. Revisiting the role of persistent neural activity during working memory. Trends in Cognitive Sciences 18, 82–89 (2014).

20. Albers, A. M., Kok, P., Toni, I., Dijkerman, H. C. & de Lange, F. P. Shared Representations for Working Memory and Mental Imagery in Early Visual Cortex. Current Biology 23, 1427–1431 (2013).

21. Iamshchinina, P. et al. Perceived and mentally rotated contents are differentially represented in cortical depth of V1. Commun Biol 4, 1069 (2021).

22. Pratte, M. S. & Tong, F. Spatial specificity of working memory representations in the early visual cortex. Journal of Vision 14, 1–12 (2014).

23. Vo, V. A. et al. Shared Representational Formats for Information Maintained in Working Memory and Information Retrieved from Long-Term Memory. Cerebral Cortex 32, 1077–1092 (2022).

24. Degutis, J. K., Weber, S., Soch, J. & Haynes, J.-D. Neural dynamics of visual working memory representation during sensory distraction. Preprint at 10.7554/eLife.99290.3 (2025).

25. Li, H.-H. & Curtis, C. E. Neural population dynamics of human working memory. Current Biology 33, 3775–3784.e4 (2023).

26. Myers, N. E. et al. Testing sensory evidence against mnemonic templates. eLife 4, e09000 (2015). 10.7554/eLife.09000.

27. Pettini, L. et al. Visual working memory representations of naturalistic images in early visual cortex are not sensory-like. bioRxiv 2025.12.11.693774 (2025) doi:10.64898/2025.12.11.693774.

28. Van Kerkoerle, T., Self, M. W. & Roelfsema, P. R. Layer-specificity in the effects of attention and working memory on activity in primary visual cortex. Nat Commun 8, 13804 (2017).

29. Pereira Seabra, J. et al. Visuospatial working memory is cortically enabled through veridical, categorical and semantic representations. bioRxiv 2025.08.08.669067 (2025) doi:10.1101/2025.08.08.669067.

30. Yan, C., Christophel, T. B., Allefeld, C. & Haynes, J.-D. Categorical working memory codes in human visual cortex. NeuroImage 274, 120149 (2023).

31. Nieder, A., Freedman, D. J. & Miller, E. K. Representation of the Quantity of Visual Items in the Primate Prefrontal Cortex. Science 297, 1708–1711 (2002).

32. Wallis, J. D., Anderson, K. C. & Miller, E. K. Single neurons in prefrontal cortex encode abstract rules. Nature 411, 953–956 (2001).

33. Nieder, A. The neuronal code for number. Nat Rev Neurosci 17, 366–382 (2016).

34. Spitzer, B., Gloel, M., Schmidt, T. T. & Blankenburg, F. Working Memory Coding of Analog Stimulus Properties in the Human Prefrontal Cortex. Cerebral Cortex 24, 2229–2236 (2014).

35. Spitzer, B. & Haegens, S. Beyond the Status Quo: A Role for Beta Oscillations in Endogenous Content (Re)Activation. eNeuro 4, ENEURO.0170-17.2017 (2017).

36. Lee, S. H., Kravitz, D. J. & Baker, C. I. Goal-dependent dissociation of visual and prefrontal cortices during working memory. Nature Neuroscience 16, 997–999 (2013).

37. Hallenbeck, G. E., Sprague, T. C., Rahmati, M., Sreenivasan, K. K. & Curtis, C. E. Working memory representations in visual cortex mediate distraction effects. Nature Communications 12, 1–18 (2021).

38. Olsson, H. & Poom, L. Visual memory needs categories. Proc. Natl. Acad. Sci. U.S.A. 102, 8776–8780 (2005).

39. Duan, Z. & Curtis, C. E. Visual working memories are abstractions of percepts. eLife 10.7554/eLife.94191.2 (2024) doi:10.7554/eLife.94191.2.

40. Libby, A. & Buschman, T. J. Rotational dynamics reduce interference between sensory and memory representations. Nat Neurosci 24, 715–726 (2021).

41. Barak, O., Tsodyks, M. & Romo, R. Neuronal population coding of parametric working memory. Journal of Neuroscience 30, 9424–9430 (2010).

42. Stokes, M. G. et al. Dynamic Coding for Cognitive Control in Prefrontal Cortex. Neuron 78, 364–375 (2013).

43. Panichello, M. F. & Buschman, T. J. Shared mechanisms underlie the control of working memory and attention. Nature 592, 601–605 (2021).

44. van Loon, A. M., Olmos-Solis, K., Fahrenfort, J. J. & Olivers, C. N. L. Current and future goals are represented in opposite patterns in object-selective cortex. eLife 7, 1–25 (2018).

45. Xu, Y. The human posterior parietal cortices orthogonalize the representation of different streams of information concurrently coded in visual working memory. PLoS Biol 22, e3002915 (2024).

46. Yu, Q., Teng, C. & Postle, B. R. Different states of priority recruit different neural representations in visual working memory. PLoS Biology 18, 1–21 (2020).

47. Fusi, S., Miller, E. K. & Rigotti, M. Why neurons mix: high dimensionality for higher cognition. Current Opinion in Neurobiology 37, 66–74 (2016).

48. Rigotti, M. et al. The importance of mixed selectivity in complex cognitive tasks. Nature 497, 585–590 (2013).

49. Bays, P. M., Schneegans, S., Ma, W. J. & Brady, T. F. Representation and computation in working memory. Nature Human Behaviour 8, 1016–1034 (2024).

50. Kiyonaga, A. & Serences, J. T. Sensory reformatting for a working visual memory. Trends in Cognitive Sciences 29, 1120–1135 (2025).

51. Linden, D. E. J., Oosterhof, N. N., Klein, C. & Downing, P. E. Mapping brain activation and information during category-specific visual working memory. Journal of Neurophysiology 107, 628–639 (2012).

52. Woodry, R., Curtis, C. E. & Winawer, J. Feedback Scales the Spatial Tuning of Cortical Responses during Both Visual Working Memory and Long-Term Memory. J. Neurosci. 45, e0681242025 (2025).

53. Zhang, M. & Yu, Q. The representation of abstract goals in working memory is supported by task-congruent neural geometry. PLoS Biol 22, e3002461 (2024).

54. Sprague, T. C., Ester, E. F. & Serences, J. T. Restoring Latent Visual Working Memory Representations in Human Cortex. Neuron 91, 694–707 (2016).

55. Mackey, W. E. & Curtis, C. E. Distinct contributions by frontal and parietal cortices support working memory. Sci Rep 7, 6188 (2017).

56. Romo, R., Brody, C. D., Hernández, A. & Lemus, L. Neuronal correlates of parametric working memory in the prefrontal cortex. Nature 399, 470–473 (1999).

57. Linde-Domingo, J. & Spitzer, B. Geometry of visuospatial working memory information in miniature gaze patterns. Nat Hum Behav 8, 336–348 (2024).

58. Kubilius, J. et al. CORnet: Modeling the Neural Mechanisms of Core Object Recognition. Preprint at 10.1101/408385 (2018).

59. Bae, G. Y. Neural evidence for categorical biases in location and orientation representations in a working memory task: EEG decoding of categorical biases. NeuroImage 240, 118366 (2021).

60. Spitzer, B. & Blankenburg, F. Stimulus-dependent EEG activity reflects internal updating of tactile working memory in humans. Proc. Natl. Acad. Sci. U.S.A. 108, 8444–8449 (2011).

61. Tam, J. & Wyble, B. Location Has a Privilege, but It Is Limited: Evidence From Probing Task-Irrelevant Location. Journal of Experimental Psychology: Learning, Memory, and Cognition 49, 1051–1067 (2023).

62. van Ede, F., Chekroud, S. R., Stokes, M. G. & Nobre, A. C. Concurrent visual and motor selection during visual working memory guided action. Nat Neurosci 22, 477–483 (2019).

63. Gayet, S., Paffen, C. L. E. & Van Der Stigchel, S. Visual Working Memory Storage Recruits Sensory Processing Areas. Trends in Cognitive Sciences 22, 189–190 (2018).

64. Teng, C. & Postle, B. R. Understanding occipital and parietal contributions to visual working memory: Commentary on Xu (2020). Visual Cognition 29, 401–408 (2021).

65. Xu, Y. Reevaluating the Sensory Account of Visual Working Memory Storage. Trends in Cognitive Sciences 21, 794–815 (2017).

66. Xu, Y. Revisit once more the sensory storage account of visual working memory. Visual Cognition 28, 433–446 (2020).

67. Wyble, B., Tam, J., Deal, I. & Bowman, H. Understanding the flexibility of working memory: Compositionality, generative processing, anchors and holistic representations. Neuroscience & Biobehavioral Reviews 179, 106387 (2025).

68. Lisman, J. E. & Idiart, M. A. Storage of 7 +/- 2 short-term memories in oscillatory subcycles. Science 267, 1512–1515 (1995).

69. Miller, E. K., Lundqvist, M. & Bastos, A. M. Working Memory 2.0. Neuron 100, 463–475 (2018).

70. Panichello, M. F. et al. Intermittent rate coding and cue-specific ensembles support working memory. Nature 636, 422–429 (2024).

71. Lundqvist, M. et al. Gamma and Beta Bursts Underlie Working Memory. Neuron 90, 152–164 (2016).

72. Jun, N. Y. et al. Coordinated multiplexing of information about separate objects in visual cortex. eLife 11, e76452 (2022).

73. Hardman, K. O., Vergauwe, E. & Ricker, T. J. Categorical working memory representations are used in delayed estimation of continuous colors. Journal of Experimental Psychology: Human Perception and Performance 43, 30–54 (2017).

74. Ricker, T. J., Souza, A. S. & Vergauwe, E. Feature identity determines representation structure in working memory. J Exp Psychol Gen 152, 2925–2940 (2023).

75. Liu, B., Nobre, A. C. & Van Ede, F. Functional but not obligatory link between microsaccades and neural modulation by covert spatial attention. Nat Commun 13, 3503 (2022).

76. Mostert, P. et al. Eye Movement-Related Confounds in Neural Decoding of Visual Working Memory Representations. eNeuro 5, ENEURO.0401-17.2018 (2018).

77. Van Ede, F., Chekroud, S. R. & Nobre, A. C. Human gaze tracks attentional focusing in memorized visual space. Nat Hum Behav 3, 462–470 (2019).

78. Bruce, C. J. & Goldberg, M. E. Primate frontal eye fields. I. Single neurons discharging before saccades. Journal of neurophysiology 53, 603–635 (1985).

79. Dell’Acqua, R. et al. ERP Evidence for Ultra-Fast Semantic Processing in the Picture–Word Interference Paradigm. Frontiers in Psychology Volume 1-2010, (2010).

80. Eimer, M. & Kiss, M. Top-down search strategies determine attentional capture in visual search: behavioral and electrophysiological evidence. Atten Percept Psychophys 72, 951–962 (2010).

81. Ehrlich, D. B. & Murray, J. D. Geometry of neural computation unifies working memory and planning. Proc. Natl. Acad. Sci. U.S.A. 119, e2115610119 (2022).

82. Nagy, D. G., Orbán, G. & Wu, C. M. Adaptive compression as a unifying framework for episodic and semantic memory. Nat Rev Psychol 4, 484–498 (2025).

83. Brodeur, M. B., Guérard, K. & Bouras, M. Bank of Standardized Stimuli (BOSS) Phase II: 930 New Normative Photos. PLOS ONE 9, e106953 (2014).

84. Esteban, O. et al. fMRIPrep 23.0.2. Software 10.5281/zenodo.852659 (2018) doi:10.5281/zenodo.852659.

85. Gorgolewski, K. et al. Nipype: a flexible, lightweight and extensible neuroimaging data processing framework in Python. Frontiers in Neuroinformatics 5, 13 (2011).

86. Gorgolewski, K. J. et al. Nipype. Software 10.5281/zenodo.596855 (2018) doi:10.5281/zenodo.596855.

87. Andersson, J. L. R., Skare, S. & Ashburner, J. How to correct susceptibility distortions in spin-echo echo-planar images: application to diffusion tensor imaging. NeuroImage 20, 870–888 (2003).

88. Avants, B. B., Epstein, C. L., Grossman, M. & Gee, J. C. Symmetric diffeomorphic image registration with cross-correlation: Evaluating automated labeling of elderly and neurodegenerative brain. Medical Image Analysis 12, 26–41 (2008).

89. Tustison, N. J. et al. N4ITK: Improved N3 Bias Correction. IEEE Transactions on Medical Imaging 29, 1310–1320 (2010).

90. Zhang, Y., Brady, M. & Smith, S. Segmentation of brain MR images through a hidden Markov random field model and the expectation-maximization algorithm. IEEE Transactions on Medical Imaging 20, 45–57 (2001).

91. Ciric, R. et al. TemplateFlow: FAIR-sharing of multi-scale, multi-species brain models. Nature Methods 19, 1568–1571 (2022).

92. Jenkinson, M., Bannister, P., Brady, M. & Smith, S. Improved Optimization for the Robust and Accurate Linear Registration and Motion Correction of Brain Images. NeuroImage 17, 825–841 (2002).

93. Cox, R. W. & Hyde, J. S. Software tools for analysis and visualization of fMRI data. NMR in Biomedicine 10, 171–178 (1997).

94. Greve, D. N. & Fischl, B. Accurate and robust brain image alignment using boundary-based registration. NeuroImage 48, 63–72 (2009).

95. Power, J. D. et al. Methods to detect, characterize, and remove motion artifact in resting state fMRI. NeuroImage 84, 320–341 (2014).

96. Satterthwaite, T. D. et al. An improved framework for confound regression and filtering for control of motion artifact in the preprocessing of resting-state functional connectivity data. NeuroImage 64, 240–256 (2013).

97. Lanczos, C. Evaluation of Noisy Data. Journal of the Society for Industrial and Applied Mathematics Series B Numerical Analysis 1, 76–85 (1964).

98. Amunts, K., Mohlberg, H., Bludau, S. & Zilles, K. Julich-Brain: A 3D probabilistic atlas of the human brain’s cytoarchitecture. Science 369, 988–992 (2020).

99. Ciric, R. et al. Benchmarking of participant-level confound regression strategies for the control of motion artifact in studies of functional connectivity. NeuroImage 154, 174–187 (2017).

100. Abraham, A. et al. Machine learning for neuroimaging with scikit-learn. Frontiers in Neuroinformatics 8, (2014).

101. Diedrichsen, J. et al. Comparing representational geometries using whitened unbiased-distance-matrix similarity. Preprint at http://arxiv.org/abs/2007.02789 (2021).

102. Walther, A. et al. Reliability of dissimilarity measures for multi-voxel pattern analysis. NeuroImage 137, 188–200 (2016).

103. Arbuckle, S. A., Pruszynski, J. A. & Diedrichsen, J. Mapping the Integration of Sensory Information across Fingers in Human Sensorimotor Cortex. J. Neurosci. 42, 5173–5185 (2022).

104. Diedrichsen, J. & Kriegeskorte, N. Representational models: A common framework for understanding encoding, pattern-component, and representational-similarity analysis. PLoS Comput Biol 13, e1005508 (2017).

105. Christophel, T. B., Iamshchinina, P., Yan, C., Allefeld, C. & Haynes, J. D. Cortical specialization for attended versus unattended working memory. Nature Neuroscience 21, 494–496 (2018).

106. RSAtoolbox Development Group. Representational Similarity Analysis 3.0.

107. Benjamini, Y. & Hochberg, Y. Controlling the False Discovery Rate: A Practical and Powerful Approach to Multiple Testing. Journal of the Royal Statistical Society: Series B (Methodological) 57, 289–300 (1995).

108. Schütt, H. H., Kipnis, A. D., Diedrichsen, J. & Kriegeskorte, N. Statistical inference on representational geometries. eLife 12, e82566 (2023).

109. Efron, B. & Tibshirani, R. An Introduction to the Bootstrap. (Chapman and Hall / CRC, New York, NY, USA, 1994).

110. Pedregosa, F. et al. Scikit-learn: Machine Learning in Python. Journal of Machine Learning Research 12, 2825–2830 (2011).

111. Yang, S., Dong, Y. & Kiyonaga, A. Flexible Working Memory in the Peripheral Nervous System. bioRxiv 10.1101/2025.09.26.678884 (2025) doi:10.1101/2025.09.26.678884.

