## Supplementary Figure for "Abstract and Concrete Working Memory Information in Human Visual Cortex"

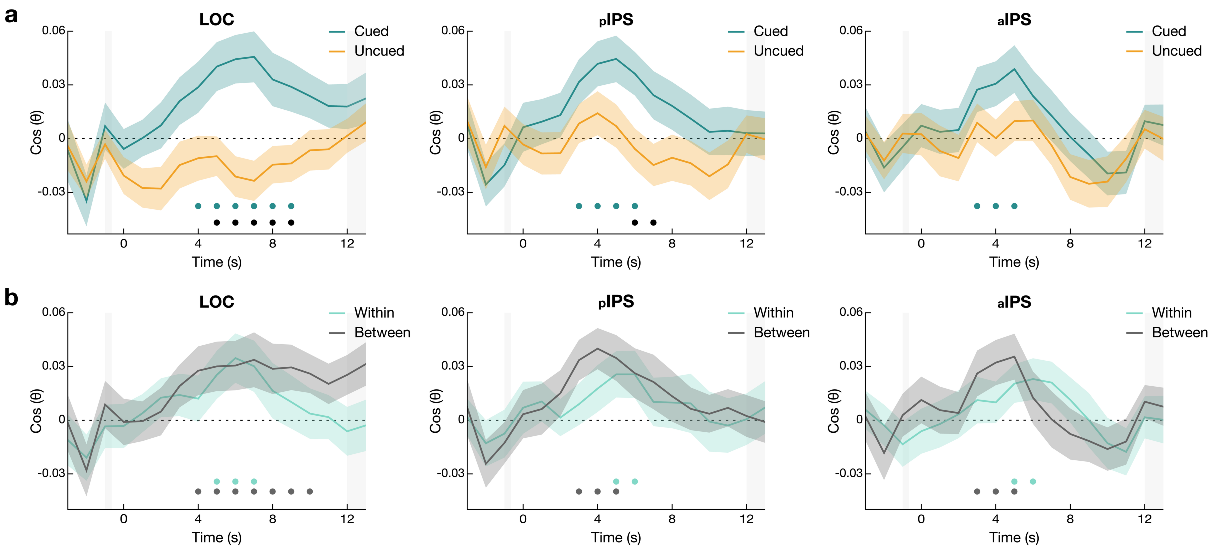


**Fig. S1 | Time course of orientation encoding (analogous to Fig. 1d and 2b) in the remaining ROIs. a,** During the WM delay, all regions of interest show significant encoding of the retro-cued (to-be-remembered) orientation. **b**, Same as *a,* for the cued orientation, but plotted separately for encoding within and between objects (see Fig. 2). All ROIs showed significant orientation encoding both within and between objects (FDR-corrected across ROIs). Vertical shadings indicate the time of the retro-cue (left) and the behavioural response period (right). Error shadings show s.e.m. The dots at the bottom indicate time points of significant ($p < 0.05$) orientation encoding compared to zero (coloured or grey) and of significant differences between conditions (black), n = 40 participants. LOC: Lateral Occipital Complex; pIPS: posterior Intraparietal Sulcus ; aIPS: anterior Intraparietal Sulcus.

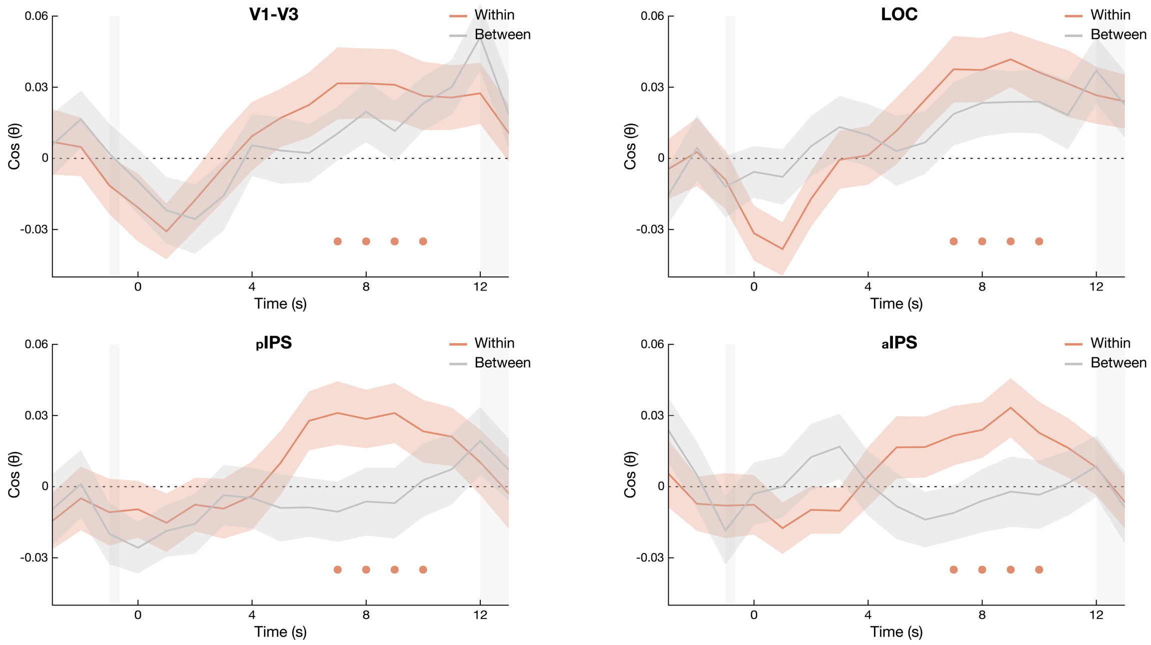


**Fig. S2 | Time course of orientation encoding with a 180° ("line") model (cf. Fig. 3b).** All regions of interest showed significant 180° orientation encoding within objects, with a peak tending towards the later part of the delay. Conversely, orientation encoding between objects did not reach significance at any time point in the delay period (see main text for delay period averages). Same conventions as Fig. S1b.

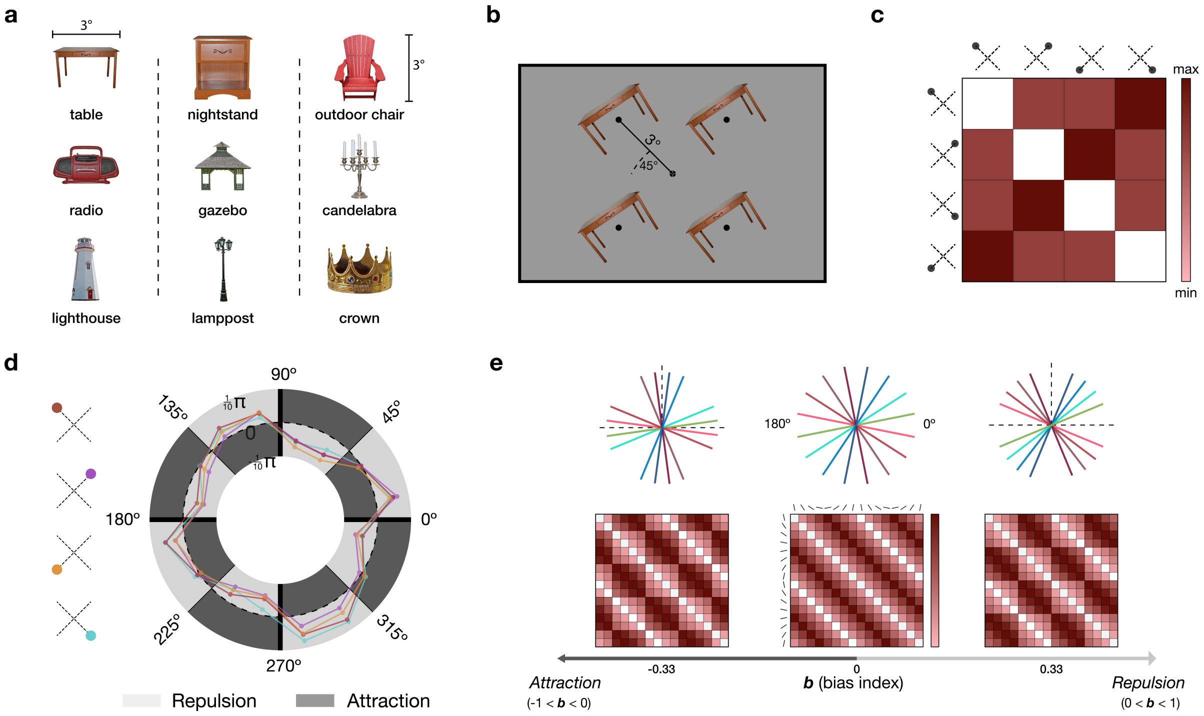


**Fig. S3 | Overview of experimental stimuli, location encoding, and "line" bias geometries. a,** We used a total of nine everyday visual objects from the BOSS database. The objects were cropped and resized to fit into a square area of 3° by 3° visual degrees. We grouped the objects into three sets, and each participant was randomly assigned to one. Each set included a combination of images with different aspect ratios (height > width, height < width, and height ≈ width). **b,** To minimise orientation-dependent gaze biases[57] while participants were asked to fixate the screen centre, we presented each sample stimulus in one of four off-centre locations (randomly chosen for each sample). The possible locations formed a square around fixation, with each location approximately 3° visual degrees distant (at a 45° angle) from the screen centre. **c**, Model RDM reflecting the pairwise Euclidean distances between the four screen locations. **d**, Polar plot of signed response error (same conventions as Fig. 4a), shown separately for each screen location (different line colours, see legend). The overall pattern of cardinal repulsion bias (see main text, Fig. 4a) was similarly evident for each of the possible screen locations. **e**, Model geometries (top) and RDMs (bottom) reflecting different levels of cardinal bias (analogous to Fig. 4b) for the 180° ("line") orientation model. Like in Fig. 4b, the bias parameter $b$ (ranging from -1 to 1) quantifies the degree of attraction to ($b < 0$) or repulsion from ($b > 0$) the cardinal axes, relative to an unbiased circular model ($\mathbf{b} = 0$).

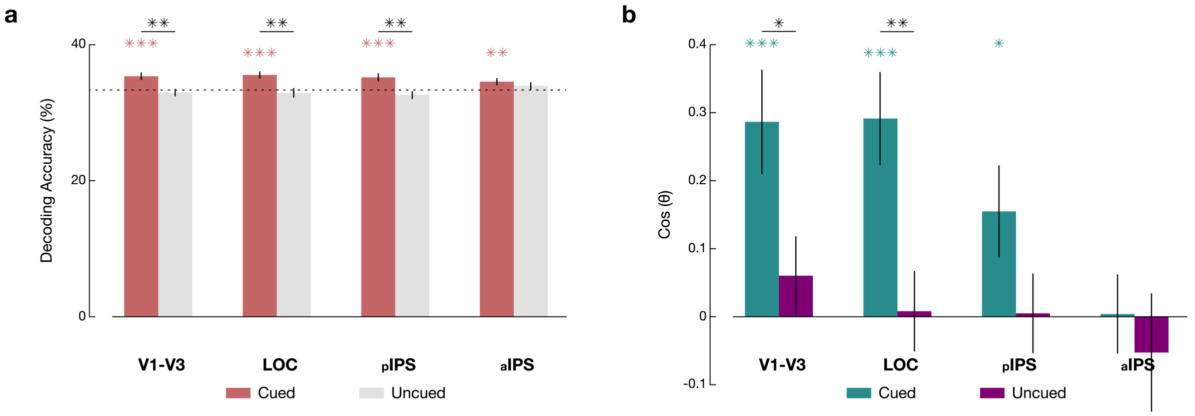


**Fig. S4 | Decoding of object identity and encoding of its spatial location. a,** Object identity decoding during the delay period (4-12 seconds after the retrocue). Decoding accuracy for the cued object was significantly above chance level (33.3%, dashed horizontal line) in all ROIs, and significantly higher than for the uncued object in all ROIs except the aIPS. See Table S7 for details. **b,** Although the samples' spatial locations were task-irrelevant (i.e., not to be reported at the WM test), we observed cue-dependent differences in location encoding in V1-V3 and LOC during the delay period, with significantly stronger location encoding for the retrocued than for the uncued sample. See Table S8 for details. Error bars show s.e.m.; coloured asterisks indicate significance compared to chance level (identity decoding) or zero (location encoding), and black asterisks indicate significance of the difference between cued and uncued samples (*$p < 0.05$, **$p < 0.01$, ***$p < 0.001$, FDR corrected). n = 40 participants. LOC: Lateral Occipital Complex; pIPS: posterior Intraparietal Sulcus; aIPS: anterior Intraparietal Sulcus.

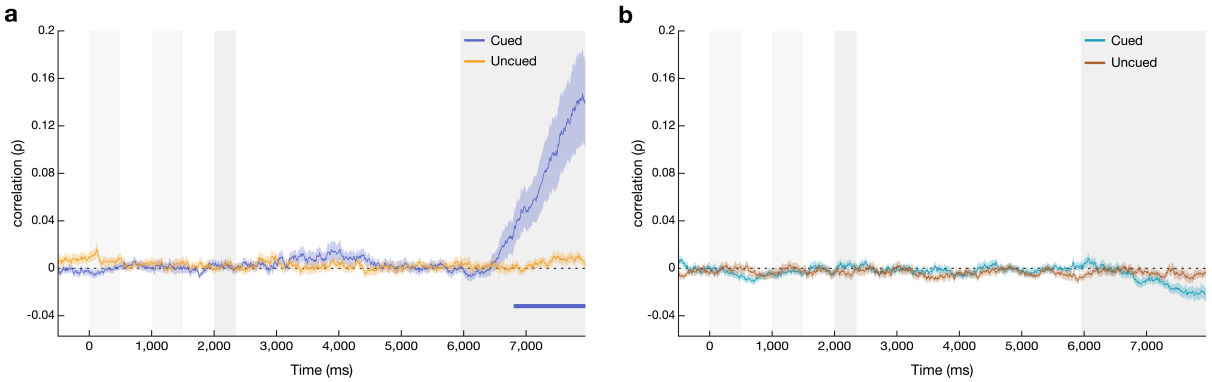


**Fig. S5 | Complementary analysis of orientation encoding in eye movements.** A previous study using a similar task setup found that when sample stimuli were presented at screen centre (i.e., at fixation), their orientations were reflected in miniature eye movements throughout the ensuing WM delay (Linde-Domingo & Spitzer, 2024). To minimise such orientation-dependent gaze patterns in our present fMRI experiment, we presented the stimuli at off-centre locations (see Fig. 1a and S4). To verify whether this modification was effective, we examined eye-tracking data from an additional WM experiment performed by a separate participant sample outside the scanner. This experiment used off-centre presentation with the same stimulus materials as the fMRI experiment, and besides a shorter delay period (3.6 seconds), closely resembled our in-scanner paradigm. We examined orientation encoding in gaze patterns using RSA analogous to our previous study[57]. Results are shown for 192 trials per participant, which matched the number of condition trials in the fMRI experiment (see *Methods*). **a,** Group-average time course of orientation encoding in gaze, using the 360° model. **b,** same as (a) but using the 180° model. Off-centre presentation effectively eliminated orientation-dependent eye movements upon sample presentation (cf.[57]). Clear 360° encoding of the cued orientation was observed only after the delay period, when participants manipulated the probe ($p_{cluster} < 0.001$, cluster-based permutation test against zero, corrected). A weak indication of 360° encoding in the delay period, approximately 2 s after the retrocue, was statistically insignificant in the fMRI-matched trial sample (no clusters with $p < 0.05$). Only when including the remaining trials (576 in total, compared to 192 in the fMRI experiment) did this encoding reach statistical significance ($p_{cluster} < 0.001$). In summary, off-centre presentation appeared effective in reducing orientation-dependent gaze dynamics to a level that is unlikely to explain our fMRI results (see Discussion). Light grey vertical markers indicate presentation times of the sample stimuli; darker grey markers indicate presentation times of the retro-cue and of the probe. Error shading shows s.e.m., n = 37 participants.

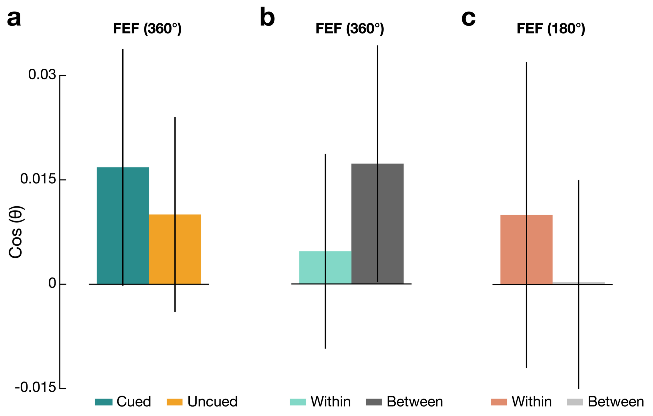


**Fig. S6 | No significant orientation encoding in FEF.** In further exploratory analysis, we examined WM-related orientation encoding in the Frontal Eye Fields (FEF; see Table S7). **a,** FEF showed no significant 360° orientation encoding for either cued or uncued orientations (both $p > 0.16$). **b,** Same as (a) for cued orientation encoding within and between objects. **c,** Same as (b) but using the 180° orientation model. Error bars show s.e.m.; $n = 40$ participants.
