## Supplementary Table for "Abstract and Concrete Working Memory Information in Human Visual Cortex"

**Supplementary Table S1.** Orientation encoding (whitened cosine similarity) with the 360° model. Detailed statistics for each ROI during the WM delay (4 -12 seconds after the retro-cue).

| ***ROI*** | ***Test*** | ***Mean* ± s.e.m.** | ***t (39)*** | ***p*** | ***BF10*** | ***Cohen's d*** |
|---|---|---|---|---|---|---|
| *V1-V3* | Cued | 0.072 ± 0.017 | 4.14 | < 0.001 | 288.89 | 0.66 |
| | Uncued | -0.014 ± 0.014 | -1.02 | 0.868 | 0.55 | 0.16 |
| | Cued - Uncued | 0.086 ± 0.019 | 4.55 | < 0.001 | 445.91 | 0.86 |
| | Within | 0.059 ± 0.016 | 3.66 | < 0.001 | 80.13 | 0.58 |
| | Between | 0.056 ± 0.018 | 3.10 | 0.005 | 19.59 | 0.49 |
| | Within - Between | 0.003 ± 0.021 | 0.15 | 0.978 | 0.17 | 0.03 |
| *LOC* | Cued | 0.057 ± 0.015 | 3.87 | < 0.001 | 136.33 | 0.61 |
| | Uncued | -0.014 ± 0.012 | -1.13 | 0.868 | 0.62 | 0.18 |
| | Cued - Uncued | 0.071 ± 0.017 | 4.15 | < 0.001 | 147.19 | 0.83 |
| | Within | 0.038 ± 0.016 | 2.34 | 0.024 | 3.82 | 0.37 |
| | Between | 0.043 ± 0.014 | 2.96 | 0.005 | 14.25 | 0.47 |
| | Within - Between | -0.005 ± 0.020 | -0.24 | 0.978 | 0.17 | 0.05 |
| *pIPS* | Cued | 0.046 ± 0.015 | 3.07 | 0.002 | 18.41 | 0.49 |
| | Uncued | -0.008 ± 0.015 | -0.54 | 0.868 | 0.39 | 0.08 |
| | Cued - Uncued | 0.054 ± 0.020 | 2.71 | 0.013 | 2.84 | 0.57 |
| | Within | 0.031 ± 0.016 | 1.94 | 0.039 | 1.85 | 0.31 |
| | Between | 0.035 ± 0.016 | 2.23 | 0.021 | 3.10 | 0.35 |
| | Within - Between | -0.004 ± 0.023 | -0.17 | 0.978 | 0.17 | 0.04 |
| *aIPS* | Cued | 0.029 ± 0.016 | 1.86 | 0.035 | 1.63 | 0.29 |
| | Uncued | -0.007 ± 0.015 | -0.44 | 0.868 | 0.37 | 0.07 |
| | Cued - Uncued | 0.035 ± 0.021 | 1.7 | 0.097 | 0.64 | 0.37 |
| | Within | 0.022 ± 0.014 | 1.61 | 0.058 | 1.12 | 0.25 |
| | Between | 0.021 ± 0.016 | 1.31 | 0.099 | 0.75 | 0.21 |
| | Within - Between | 0.001 ± 0.019 | 0.03 | 0.978 | 0.17 | 0.01 |

**Supplementary Table S2.** Results from 2 x 2 repeated-measures ANOVAs with the factors ROI (left column) and type of orientation encoding (within vs between objects).

| *ROI* | *Test* | *MS* | *F (1,39)* | *p* |
|---|---|---|---|---|
| *V1-V3 / LOC* | WI-BW | 0.01 | 1.82 | 0.185 |
| | ROI | < 0.01 | < 0.01 | 0.963 |
| | ROI x WI-BW | < 0.01 | 0.23 | 0.635 |
| *V1-V3 / pIPS* | WI-BW | <0.01 | < 0.01 | 0.987 |
| | ROI | 0.02 | 3.84 | 0.057 |
| | ROI x WI-BW | < 0.01 | 0.09 | 0.765 |
| *V1-V3 / aIPS* | WI-BW | < 0.01 | 0.01 | 0.906 |
| | ROI | 0.05 | 5.31 | 0.027 |
| | ROI x WI-BW. | < 0.01 | 0.01 | 0.915 |
| *LOC / pIPS* | WI-BW | < 0.01 | 0.05 | 0.824 |
| | ROI | < 0.01 | 0.50 | 0.484 |
| | ROI x WI-BW | < 0.01 | < 0.01 | 0.948 |
| *LOC / aIPS* | WI-BW | < 0.01 | < 0.01 | 0.987 |
| | ROI | 0.02 | 3.84 | 0.057 |
| | ROI x WI-BW | < 0.01 | 0.09 | 0.52 |
| *pIPS / aIPS* | WI-BW | < 0.01 | 0.02 | 0.898 |
| | ROI | 0.01 | 2.43 | 0.127 |
| | ROI x WI-BW | < 0.01 | 0.07 | 0.797 |

**Supplementary Table S3.** Orientation encoding with the 180° ("line ") model. Detailed statistics for each ROI during the WM delay (4 -12 seconds after the retro-cue).

| ***ROI*** | ***Test*** | ***Mean* ± s.e.m.** | ***t (39)*** | ***p*** | ***BF10*** | ***Cohen's d*** |
|---|---|---|---|---|---|---|
| *V1-V3* | Within | 0.037 ± 0.015 | 2.46 | 0.009 | 4.86 | 0.39 |
| | Between | 0.019 ± 0.014 | 1.40 | 0.265 | 0.84 | 0.22 |
| | Within - Between | 0.018 ± 0.019 | 0.96 | 0.341 | 0.26 | 0.20 |
| *LOC* | Within | 0.045 ± 0.015 | 3.03 | 0.008 | 16.72 | 0.48 |
| | Between | 0.020 ± 0.018 | 1.13 | 0.265 | 0.62 | 0.18 |
| | Within - Between | 0.025 ± 0.024 | 1.06 | 0.341 | 0.29 | 0.24 |
| *pIPS* | Within | 0.031 ± 0.013 | 2.43 | 0.009 | 4.6 | 0.38 |
| | Between | -0.005 ± 0.016 | -0.33 | 0.724 | 0.36 | 0.05 |
| | Within - Between | 0.036 ± 0.018 | 1.95 | 0.116 | 0.95 | 0.39 |
| *aIPS* | Within | 0.034 ± 0.013 | 2.57 | 0.009 | 6.13 | 0.41 |
| | Between | -0.009 ± 0.015 | -0.60 | 0.724 | 0.40 | 0.10 |
| | Within - Between | 0.043 ± 0.018 | 2.66 | 0.045 | 1.86 | 0.48 |

**Supplementary Table S4.** Similarities of CORnet-S patterns at each pooling layer to our 360° and 180° orientation models (measured in whitened correlation), respectively (see Methods for bootstrapping analysis and confidence interval calculation).

| | | **360°** | | **180°** | |
|---|---|---|---|---|---|
| **Layer** | **Test** | **Mean (r)** | **p** | **Mean (r)** | **p** |
| *I (V1)* | Within | 0.048 | 0.238 | 0.586 | < 0.001 |
| | Between | 0.004 | 0.491 | -0.258 | > 0.999 |
| | WI - BW | 0.446 | 0.566 | 0.843 | < 0.001 |
| *I I (V2)* | Within | -0.020 | 0.578 | 0.607 | < 0.001 |
| | Between | -0.022 | 0.795 | -0.262 | > 0.999 |
| | WI - BW | 0.002 | 0.877 | 0.869 | < 0.001 |
| *I I I (V4)* | Within | 0.179 | 0.019 | 0.622 | < 0.001 |
| | Between | -0.006 | 0.660 | -0.227 | > 0.999 |
| | WI - BW | 0.186 | 0.089 | 0.848 | < 0.001 |
| *IV (IT)* | Within | 0.583 | < 0.001 | 0.486 | < 0.001 |
| | Between | -0.046 | 0.999 | -0.036 | > 0.999 |
| | WI-BW | 0.629 | < 0.001 | 0.522 | < 0.001 |

**Supplementary Table S5.** Estimates of bias parameter *b* (*b* < 0: attraction, *b* > 0: repulsion; see Methods and Results) from fitting the parameterised 360° ("circle") model—inferential and Bayesian statistics (two-tailed) against zero.

| | **b (mean ± s.e.m.)** | **t (39)** | **p** | **BF10** | **Cohen's d** |
|---|---|---|---|---|---|
| *Behavioral* | 0.375 ± 0.022 | 16.82 | < 0.001 | >1000 | 2.66 |
| *V1-V3* | 0.009 ± 0.129 | 0.07 | 0.939 | 0.17 | 0.01 |
| *LOC* | 0.028 ± 0.129 | 0.22 | 0.939 | 0.17 | 0.03 |
| *pIPS* | 0.098 ± 0.137 | 0.72 | 0.792 | 0.22 | 0.11 |
| *aIPS* | 0.396 ± 0.126 | 3.15 | 0.008 | 11.10 | 0.50 |

**Supplementary Table S6.** Same as Table S5, but for the 180° (line) model (within-object encoding; see Fig. 3b).

| | b (mean ± s.e.m.) | t (39) | p | BF10 | Cohen's d |
|---|---|---|---|---|---|
| *Behavioral* | 0.004 ± 0.031 | 0.14 | 0.892 | 0.17 | 0.02 |
| *V1-V3* | 0.044 ± 0.126 | 0.348 | 0.911 | 0.18 | 0.06 |
| *LOC* | -0.003 ± 0.123 | -0.02 | 0.981 | 0.17 | < 0.01 |
| *pIPS* | 0.049 ± 0.133 | 0.37 | 0.911 | 0.18 | 0.06 |
| *aIPS* | -0.143 ± 0.125 | -1.14 | 0.650 | 0.31 | 0.18 |

**Supplementary Table S7.** Object identity decoding accuracy (% correct classification) for cued and uncued samples during the WM delay (4 -12 seconds after the retro-cue). The cued and uncued results t-tests are against a chance level of 33.33% (one-tailed t-test); all other t-tests are two-tailed.

| *ROI* | *Test* | *Mean* ± s.e.m. | *t (39)* | *p* |
|---|---|---|---|---|
| *V1-V3* | Cued | 35.36 ± 0.55 | 3.68 | < 0.001 |
| | Uncued | 32.95 ± 0.54 | -0.71 | 0.909 |
| | Cued - Uncued | 2.40 ± 0.77 | 3.14 | 0.005 |
| *LOC* | Cued | 35.55 ± 0.54 | 4.10 | < 0.001 |
| | Uncued | 32.92 ± 0.69 | -0.59 | 0.909 |
| | Cued - Uncued | 2.68 ± 0.84 | 3.11 | 0.005 |
| *pIPS* | Cued | 35.27 ± 0.51 | 3.81 | < 0.001 |
| | Uncued | 32.63 ± 0.52 | -1.36 | 0.909 |
| | Cued - Uncued | 2.65 ± 0.67 | 3.97 | 0.001 |
| *aIPS* | Cued | 34.73 ± 0.51 | 2.72 | 0.005 |
| | Uncued | 33.89 ± 0.56 | 0.98 | 0.667 |
| | Cued - Uncued | 0.85 ± 0.72 | 1.16 | 0.252 |

**Supplementary Table S8.** Encoding of the samples' screen location. Detailed statistics for each ROI during the WM delay (4 -12 seconds after the retro-cue).

| ROI | Test | Mean ± s.e.m. | t (39) | p | BF10 | Cohen's D |
|---|---|---|---|---|---|---|
| *V1-V3* | Cued | 0.286 ± 0.076 | 3.73 | < 0.001 | 94.94 | 0.59 |
| | Uncued | 0.060 ± 0.058 | 1.04 | 0.610 | 0.56 | 0.16 |
| | Cued - Uncued | 0.226 ± 0.089 | 2.55 | 0.030 | 2.91 | 0.53 |
| *LOC* | Cued | 0.291 ± 0.068 | 4.26 | < 0.001 | 397.91 | 0.67 |
| | Uncued | 0.008 ± 0.059 | 0.14 | 0.662 | 0.34 | 0.02 |
| | Cued - Uncued | 0.283 ± 0.083 | 3.40 | 0.008 | 20.54 | 0.70 |
| *pIPS* | Cued | 0.155 ± 0.065 | 2.37 | 0.015 | 4.10 | 0.38 |
| | Uncued | 0.001 ± 0.058 | 0.01 | 0.662 | 0.34 | 0.01 |
| | Cued - Uncued | 0.154 ± 0.088 | 1.75 | 0.117 | 0.68 | 0.39 |
| *aIPS* | Cued | 0.002 ± 0.068 | 0.03 | 0.489 | 0.34 | 0.01 |
| | Uncued | -0.051 ± 0.061 | -0.83 | 0.795 | 0.47 | 0.13 |
| | Cued - Uncued | 0.053 ± 0.087 | 0.61 | 0.548 | 0.20 | 0.13 |

**Supplementary Table S9.** Regions of Interest (ROI) derived from the Julich-Brain atlas 3.0 and the anatomical loci that comprise them

| ROI | Description | Anatomical loci |
|---|---|---|
| *V1-V3* | Early Visual Cortex | hOc1, hOc2, hOc3d, hOc3v |
| *LOC* | Lateral Occipital Cortex | hOc4lp, hOc4la, hOc5 |
| *pIPS* | Posterior intraparietal sulcus | hIP4, hIP5, hIP7, hIP8 |
| *aIPS* | Anterior intraparietal sulcus | hIP1, hIP2, hIP3, hIP6 |
| *FEF* | Frontal Eye Fields | 6d1 |
